# The metabolome and lipidome of colorectal adenomas and cancers

**DOI:** 10.1101/2021.06.01.446510

**Authors:** Endre Laczko, Christine Manser, Giancarlo Marra

**Affiliations:** Functional Genomics Center of the University of Zurich and ETH Zurich, Switzerland; Division of Gastroenterology and Hepatology, University Hospital Zurich, Switzerland; Institute of Molecular Cancer Research, University of Zurich, Switzerland; Department of Gastroenterology and Hepatology, Cantonal Hospital of Frauenfeld, Switzerland

**Keywords:** Colorectal adenoma, Colorectal cancer, Metabolome, Lipidome, Liquid chromatography-mass spectrometry

## Abstract

**Introduction:** In-depth knowledge of metabolic dysregulations in colorectal cancer (CRC) (and other cancers as well) is essential for developing treatments that specifically kill neoplastic cells. It may also allow us to pinpoint metabolites or lipids with potential for development as tumor biomarkers for use in body-fluid or breath assays. CRC onset is preceded by an interval of ∼10 years characterized by the presence of precancerous lesions, and our previous studies have revealed epigenomic, transcriptomic, and proteomic evidence in these lesions of certain metabolic changes typical of CRC. These findings prompted us to conduct untargeted metabolomic and lipidomic analyses of CRCs and colorectal adenomas (the most common precancerous lesions of the gut).

**Methods:** We analyzed 29 endoscopically collected tumor tissue samples (29 adenomas [ADNs], 10 CRCs, each with a colon segment-matched sample of normal mucosa [i.e., 29 NM-ADN, 10 NM-CRC]). The freshly collected samples were promptly frozen in liquid nitrogen and later processed to obtain metabolite and lipid extracts. Each of the 78 samples was analyzed with nano-flow LC-MS/MS (liquid chromatography with mass spectrometry) to characterize its metabolome (using HILIC, Hydrophilic Interaction Liquid Chromatography) and lipidome (using RP, Reversed Phase chromatography). The data acquired were processed using Progenesis QI. For statistical and multivariate analysis of the resulting peak tables, we used basic R packages and the R package made4.

**Results:** Unsupervised between-group analysis based on the full set of detected metabolites (n=1830) and lipids (n=2365) clearly discriminated ADNs and CRCs from their matched samples of normal mucosa at both the metabolome and lipidome levels. Compared with the NM-ADN, the ADNs contained significantly different levels of 14.6% of the metabolites and 10.8% of the lipids. Fewer compounds (9.1% of metabolites, 6.2% of lipids) displayed differential abundance in CRCs (vs. NM-CRC). The metabolome and lipidome of the NM-ADN also differed from those of the NM-CRC, probably reflecting the presence of a field cancerization effect exerted by the invasive tumors. A substantial number of metabolites (n=340) and lipids (n=201) also displayed abundance differentials across the sequential tumorigenic stages represented by the NM-ADN (considered more representative of NM from a lesion-free colon) → ADN → CRC. In most cases, the trend consisted of progressive increases or progressive decreases in abundance as the tumorigenesis advanced.

**Conclusions:** Our findings provide a preliminary picture of the progressive metabolomic and lipidomic changes occurring during the adenomatous phase of colorectal tumorigenesis. Once definitively annotated, the numerous differentially abundant compounds detected in this study may well shed valuable light on the metabolic dysregulations occurring during this process and provide useful clues for the development of novel tools for the diagnosis and treatment of colorectal tumors.

## Introduction

Colorectal cancer (CRC) is frequent in the Western world. In Switzerland, for example, it is the second most common cancer diagnosed in women (after breast cancer) and the third most common in men (after cancers of the prostate and lung) ^1^. CRC is, however, singularly preventable: its onset is preceded by an interval of approximately 10-15 years, during which benign CRC-precursor lesions are detectable in the large intestine ^2–5^. This broad time window can be exploited for the prevention and early diagnosis of CRC via colonoscopy-based screening, with prompt endoscopic resection of suspicious lesions. Use of this approach has been largely responsible for the declines in CRC mortality and incidence rates observed over the past 40 years ^6,7^.

Sixty percent of all precancerous colorectal lesions are conventional adenomas (ADN) ^8,9^. The CRC risk associated with these genuinely dysplastic tumors has been reliably estimated thanks to decades of screening colonoscopy. Increases in size are accompanied by increases in the cellular and architectural dysplasia of the adenoma and also in its cancer risk^10–12^. The remaining 40% of endoscopically identified benign colorectal lesions are serrated tumors. They are generally non-dysplastic, and while the cancer risk of some serrated tumors (i.e., hyperplastic polyps) ranges from absent to very low, the risks associated with large sessile serrated lesions in the proximal colon and traditional serrated adenomas are similar to that of conventional adenomas ^9,13^.

The *normal mucosa - to - ADN - to - CRC* transformation process was molecularly characterized for the first time in the late eighties by Vogelstein et al. The process was linked to a temporal sequence of genetic alterations^14^, which were later more precisely localized in the *APC, KRAS, SMAD4*, and *TP53* genes and confirmed as “driver” mutations with the advent of deep sequencing ^15,16^. Indeed, the signaling pathways displaying the most marked dysregulation in colorectal tumors are those in which the products of these few genes play pivotal roles^16^. Mutation of the *Adenomatous polyposis coli* gene (*APC*) is one of the earliest genetic alterations in the adenomatous tumorigenic process. It is the most frequent mutational driver in ADNs ^17^ and consequently one of the most commonly identified mutations in the CRCs that develop along the adenomatous pathway^15,16,18^. The APC protein’s involvement in the degradation of beta-catenin makes it a key intracellular transducer in the canonical Wnt signaling pathway. Its structural alteration or absence therefore causes aberrant activation of this pathway, which is normally active exclusively at the base of the normal colorectal crypts where stem cells reside ^19^.

Our previous transcriptomic characterization of the Wnt signaling dysregulation that occurs in ADNs and CRCs ^20– 24^ revealed many other alterations in the precancerous lesions that were unrelated to the Wnt pathway, which prompted further investigation at the epigenomic and proteomic levels. More recently, we have also explored the relationship between gene expression and epigenetic changes in ADNs and CRCs and discovered that the dramatic changes in DNA methylation found in the cancers—which are variably associated with transcriptional outcomes—are already evident (albeit in milder forms) in their precursors ^25,26^. Our proteomic studies have also revealed important correlations between transcript and protein levels in ADNs as well as in CRCs ^27,28^. All of these findings have been complemented by the results of the study presented here, in which we explored the metabolomic and lipidomic profiles of the normal mucosa of the large intestine, colorectal ADNs, and the CRCs that arise from the latter lesions.

## Methods

### Human tissue samples

Colorectal tissues were prospectively collected from patients undergoing colonoscopy in the Gastroenterology Unit of the University Hospital in Zurich, Switzerland. The protocol was approved by the local ethics committee, and each donor provided written informed consent to sample collection, data analysis, and publication of the findings. The series comprised 39 colorectal tumors from a total of 31 patients: 29 precancerous lesions (polypoid or nonpolypoid conventional adenomas [ADNs]) and 10 CRCs ^29^ (**Table 1**). All tumors had maximum diameters of at least 10 mm (to ensure that our sampling procedure left adequate tissue for the histological examination). For each tumor, we also collected a colon segment-matched sample of normal mucosa (NM) 2 to 5 cm from the lesion (referred to hereafter as NM-ADN and NM-CRC). The 78 tissue samples thus obtained were subjected to metabolome and lipidome analyses.

**Table 1.** Characteristics of the patients with ADNs and CRC analyzed.

| Table 1. Characteristics of the patients with ADNs and CRC analyzed. |  |  |  |  |  |  |  |  |  |
| --- | --- | --- | --- | --- | --- | --- | --- | --- | --- |
| Tumor no. * | Pt. Age | Pt. Sex | Location in the colon | Max. lesion diameter (mm) | Paris class. # | Microscopic appearance | Tumor stage * | No. of lesions at study colonoscopy | No. of previously excised lesions ‡ |
| **ADNs** |  |  |  |  |  |  |  |  |  |
| AA45 | 65 | M | C | 10 | Ila / I Ib | TVA | HGD | 3 | 0 |
| AH49 | 61 | M | A | 10 | Is | TA | LGD | 3 | 0 |
| AP53 * | 57 | M | A | 40 | Is | TA | LGD | 5 | 2 |
| BR55 ** | 55 | M | A | 12 | Ip | TA | LGD | 3 | 0 |
| BR553 ** | 55 | M | S | 25 | Ip | TVA | HGD | 3 | 0 |
| EK43 | 67 | M | S | 10 | Ila | TVA | HGD | 4 | 0 |
| EM34 | 76 | F | C | 18 | I Ib / I Ic | TVA | HGD | 1 | 0 |
| FA57 | 53 | M | SF | 30 | Ip | TA | HGD | 1 | 0 |
| FK49 | 61 | M | C | 25 | Is | TVA | LGD | 0 | 0 |
| FZ67 | 43 | M | S | 26 | Ip | TA | HGD | 1 | 0 |
| GM43 | 67 | M | T | 12 | Is | TVA | HGD | 2 | 0 |
| HJ41 | 69 | M | S | 20 | Ip | TVA | HGD | 6 | 0 |
| HM52 | 57 | M | S | 20 | Ip | TA | HGD | 1 | 0 |
| HS36 *** | 74 | M | HF | 15 | Ila | TA | HGD | 3 | 0 |
| HS362 *** | 74 | M | S | 30 | Is | TVA | HGD | 3 | 0 |
| HT54 | 56 | M | S | 27 | Is | TA | HGD | 1 | 0 |
| IR57 | 53 | M | S | 10 | Is | TA | HGD | 3 | 0 |
| KE55 | 55 | M | C | 40 | Ila / I Ib | TVA | LGD | 1 | 0 |
| KS37 **** | 73 | M | C | 25 | Is | TA | LGD | 4 | 0 |
| KS371 **** | 73 | M | A | 30 | Ip | TA | LGD | 4 | 0 |
| KS47 | 63 | M | T | 10 | Ila | TA | LGD | 4 | 0 |
| LF35 | 75 | M | R | 12 | Is | TVA | LGD | 0 | 6 |
| MK37 | 72 | M | S | 20 | Ip | TVA | LGD | 2 | 0 |
| OM37 | 73 | M | HF | 20 | Is | TVA | HGD | 4 | 3 |
| PH36 | 74 | M | D | 15 | Is | TVA | LGD | 6 | 3 |
| SR33 ***** | 77 | F | C | 30 | Ila | TVA | HGD | 5 | 0 |
| SR332 ***** | 77 | F | A | 30 | Ila | TA | HGD | 5 | 0 |
| SZ33 | 77 | M | HF | 10 | Ip | TA | HGD | 6 | 2 |
| WA52 | 58 | M | S | 25 | Ip | TA | LGD | 0 | 0 |
| **CRCs** |  |  |  |  |  |  |  |  |  |
| AP531 * | 57 | M | D | 30 | cancer | cancer | pT1, pN1 | 5 | 2 |
| AP532 * | 57 | M | S | 40 | cancer | cancer | pT1, pN1 | 5 | 2 |
| GA39 | 71 | M | HF | 40 | cancer | cancer | pT2, pN0 | 6 | 0 |
| HH53 | 57 | M | S | 23 | cancer | cancer | pTis | 4 | 1 |
| HS361 *** | 74 | M | S | 40 | cancer | cancer | pT1, pN0 | 3 | 0 |
| HW38 | 72 | M | A | 30 | cancer | cancer | pT3, pN0 | 1 | 0 |
| RH61 | 48 | F | S | 40 | cancer | cancer | pTis | 1 | 0 |
| SR331 ***** | 77 | F | C | 30 | cancer | cancer | pTis | 5 | 0 |
| TD | 75 | F | C | 30 | cancer | cancer | pT4, pN2, cM1 | na | na |
| TO | 62 | M | S | 40 | cancer | cancer | pT3, pN1 | na | na |
| *, **, ***, ****, *****: indicate multiple tumors from the same patient (e.g., ADN AP53 and CRCs AP531 and AP532). |  |  |  |  |  |  |  |  |  |
| # Macroscopic appearance of neoplastic lesions was classified according to Paris Endoscopic Classification: Update on the Paris classification of superficial neoplastic lesions in the digestive tract; Endoscopy 37, 570-578 (2005) |  |  |  |  |  |  |  |  |  |
| * For ADNs: Low- or high-grade dysplasia; for CRCs, tumor stage as defined by the WHO classification of tumors of the digestive system, editorial and consensus conference in Lyon, France, November 6-9, 1999. IARC |  |  |  |  |  |  |  |  |  |
| ‡ Total no. adenomas detected and excised during previous colonoscopies. |  |  |  |  |  |  |  |  |  |
| Abbreviations: ADN, conventional adenoma; CRC, colorectal carcinoma; M, male; F, female; C, cecum; A, ascending colon; HF, hepatic flexure; SF, splenic flexure; T, transverse colon; D, descending colon; Pt, patient; S, sigmoid colon; R, rectum; TA, tubular adenoma; TVA, tubulovillous adenoma; VA, villous adenoma; LGD, low-grade dysplasia; HGD, high-grade dysplasia; Tis: carcinoma in situ. |  |  |  |  |  |  |  |  |  |

### Extraction and analysis of metabolites and lipids

Immediately after collection, each tissue biopsy was frozen in liquid nitrogen and stored at -80°C. The fresh weight of each frozen specimen was determined, and extraction solvent was added to the sample at a ratio of 100 μl per 3 mg tissue. The extraction solvent consisted of methanol (98 parts), internal metabolite standards solution (1 part), and internal lipid standards solution (1 part) (v/v, composition of internal standards mixture in **Supplementary Table 1**). Immediately after solvent addition, the tissue was mechanically disrupted in a glass tissue grinder (Potter-Elvehjem homogenizer from Wheaton, Millville, New Jersey, USA) on ice. The homogenate was centrifuged for 5 minutes at 4000 *g*, and the clear methanol extract was collected for further analysis.

Metabolome and lipidome analyses of the tissue extracts were carried out with two different methods, each based on capillary liquid chromatography coupled, via a nano ESI (electro-spray ionization) source, to a hybrid tandem mass spectrometer (Synapt G2 HDMS, Waters, Manchester, UK), and both using self-packed capillary spray tip columns. We used HILIC (hydrophilic interaction liquid chromatography) at basic conditions (pH 9.0) for the metabolome analysis and RP (Reversed Phase) chromatography for the lipidome analysis (**Supplementary Table 2)**.

For the metabolome analysis, an aliquot of the methanol extract was vacuum-dried and then re-dissolved in ultra-pure water. This aqueous solution was diluted five times with the injection solvent mixture consisting of 50 mM ammonium acetate, 90% acetonitrile, 9% methanol, and 1% ultrapure water adjusted to pH 9.0 with concentrated ammonia hydroxide solution. Before undergoing Liquid Chromatography - Mass Spectrometry (LC-MS) analysis, the processed samples were centrifuged for 2 minutes at 4000 *g* to remove residual precipitates. For the lipidome analysis, we diluted a second aliquot of the methanol extract five times with solvent A used for the RP method.

Mass spectrometry (MS) was performed using negative (metabolites) or positive (lipids) ESI polarization at a rate of 0.3 scans per second (3Hz) in MS^E^ mode (all ion fragmentation mode MS/MS) ^30^. In negative mode, the MS parameters were: 1.2 kV capillary voltage, 30 V sampling cone voltage, and 3 V extraction cone voltage. The source temperature was set to 100°C, and sheet gas (nitrogen) was applied. For lipids and hydrophobic metabolites, we used positive-mode acquisition with the same settings used in negative mode with the exception of kV capillary voltage, which was increased to 3.0. Trap Collision Energy was set to a 20-40 V ramp. Mass analysis was set to resolution mode (FWHM ∼20000 at m/z 554 or 556, for ESI- or ESI+ respectively), and an m/z scn range of 50-1200 Th was selected.

### LC-MS^E^ data processing

Continuum LC-MS^e^ data were processed using Progenesis QI 3.0 software (Nonlinear Dynamics, A Waters Company, Newcastle upon Tyne, UK). The resulting peak intensities were normalized to the median intensity of the internal standards (Supplementary Table 1) and to all ions detected (“All ions mode” in Progenesis QI). Only data subjected to the latter normalization procedure was used for further analysis. Matches with a 0.01-Da tolerance to the Human Metabolome Database (www.hmdb.ca) and LipidMaps (www.lipidmaps.org) were used for tentative annotation of the identified m/z ions. See **Table 2** for an overview of the data processing procedure and parameter settings. We also matched the observed m/z and retention-time values to those of a custom-built library of 190 well-characterized, reliably detectable and quantifiable core metabolites, which were previously analyzed under identical conditions on the same instrument (**Supplementary Table 3**).

**Table 2.** Summary of the Progenesis QI*-mediated capLC-nanoESI-MS/MS data processing procedure and annotation grades of the detected ions.

|  | Metabolites | Lipids ESI+ | Lipids ESI- |
| --- | --- | --- | --- |
| Compounds | 1830 | 1984 | 381 |
| Matches to custom library (identified) | **62** |  |  |
| Matches to databases: |  |  |  |
| HMDB ** | **709** | - | - |
| LipidM&B ** | - | **889** | **193** |
| ID annotation grade 4.5 - 5 *** | 24 | 24 | 1 |
| ID annotation grade 3.5 - 4 *** | 79 | 139 | 14 |
| ID annotation grade 1 - 3 *** | 606 | 726 | 178 |

Abbreviations: ESI, Electron Spray Ionization; HMDB, Human Metabolome Database; ID, identification.

| * Progenesis QI settings |  |  |  |
| --- | --- | --- | --- |
|  | Metabolites | Lipids ESI+ | Lipids ESI- |
| Ion charges | -1, -2 | 1 | -1, -2 |
| Adducts | M-H, M+Cl,<br>M-H<sub>2</sub>O-H,<br>M+Na-2H,<br>M-2H | M+H, 2M+H,<br>M+Na, M+NH<sub>4</sub>,<br>M-H<sub>2</sub>O+H | M-H, M+Cl,<br>M-H<sub>2</sub>O-H,<br>M+Na-2H,<br>M-2H |
| Retention Time range (min) | 3.6 - 9.5 | 5.5 - 14.8 | 4.0 - 14.0 |
| Sensitivity setting | 3 | 3 | 5 |

\*\* Databases used for annotation. Database matches within a tolerance of 5mD were accepted as tentative identifications. Multiple annotations for a single compound were ranked according to scores that weigh the deviation from the theoretical mass, the theoretical natural isotopolog composition, and the theoretical fragmentation spectra, as well as the quality of the neutral mass calculation (no. of detected adducts). Supplementary Tables 5 and 6 shows the detected compound ions, all possible annotations with scores, and the list of the compounds with top-ranked unique identifications.
\*\*\* The highest ranked annotation in accordance with Schrimpe-Rutledge et al, 2016, JAMS 27: 1897-1905. Briefly, the ID grade is the sum of 5 scores, each assigned to a feature indicative of the quality of the database-derived m/z annotation. **Score 1.** For a given compound, a pattern of adducts was observed and a neutral mass was derived: TRUE (1 point), FALSE (0 points). **Score 2.** Within the search tolerance used for the database query, a matching annotation was found: TRUE (1 point), FALSE (0 point). **Score 3.** At least one theoretical fragment was observed: TRUE (1 point), FALSE (0 points). **Score 4.** The difference between the observed neutral mass or m/z was: less than 10ppm (1 point); less than or equal to 25ppm (0.5 points); greater than 25ppm (0 points). **Score 5.** The similarity between the observed isotopologue pattern and the theoretical pattern of the data base match is: at least 98% (1 point); less than 98% but greater than or equal to 90% (0.5 point); less than 90% (0 points).

### Between-group analysis (BGA)

Omics datasets are typically characterized by high dimensionality and a markedly larger number of variables (i.e., compounds, such as metabolites and lipids) than of samples (e.g., tissue samples). A situation of this type fails to satisfy the requirements for many commonly used statistical approaches (e.g., univariate tests, analysis of variance, and supervised classification techniques). In these cases, unsupervised multivariate methods (i.e., ordination techniques), such as principal components analysis, are often used to reduce the dimensionality of the data set by replacing the original hundreds or thousands of variables with a small number of inferred (latent) variables, i.e., principal components. Principal components retain as much as possible of the variance and data structure of the original high-dimensional data set. However, unsupervised ordination techniques do not consider grouping of samples, so they are not ideal for analyzing and visualizing differences between groups and for finding optimal classifiers, i.e., markers discriminating between groups like normal colon, ADNs, and CRCs. Culhane et al. adapted a classification method used with ecology data, which relies on ordination techniques to circumvent the problem of excess variables and incorporates group information by ordinating the data based on predefined groups ^31^. This method, known as between group analysis (BGA), ordinates or projects the samples onto a reduced space of N – 1 dimensions (i.e., BGA axes), where *N* is the total number of defined groups. The BGA axes are set to discriminate each group optimally from other groups. In other words, the ordination procedure is defined using only the group centers and then applied for the ordination of the individual samples.

The BGA function is available as part of the R package *made4* ^32^. We performed BGA to group our tissue samples based on their histology. BGA allowed us to investigate and visualize associations between these histologic phenotypes and changes during tumorigenesis in the pool sizes of metabolites and lipids. Our classification approach therefore assigned each tissue sample and compound to one of the three histologic phenotypes (NM, ADNs, CRCs).

Volcano plots, BGA plots, and heatmaps were generated using the basic plotting functions of R and the plotting functions of the R package *made4*. P-values were calculated on log10(x+1) transformed data, and fold changes are reported as log2 fold changes.

The raw data generated and analyzed during the current study are available in the *MetaboLights* repository (accession numbers MTBLS2202).

## Results

We used capLC-nanoESI-MS/MS for an untargeted analysis of the metabolomes and lipidomes of NM and both precancerous and invasive tumors of the large intestine (ADNs and CRCs). We chose two chromatographic setups, HILIC and RP, to measure a broad range of polar and non-polar compounds in these tissues (see *Methods*). The features of tissue samples are shown in **Table 1** and **Supplementary Table 4**; metabolite- and lipid-intensity tables are available in **Supplementary Tables 5** and **6**, respectively. A total of 1830 metabolites and 2365 lipids were detected. The compound identification procedure yielded a unique annotation for 760 (41.5%) of the metabolites and 1028 (43.5%) of the lipids; 102 metabolites (13.4%) and 146 lipids (14.2%) were identified with certainty (identity grade 4.5 - 5) or a high probability (identity grade 3.5 - 4) on the basis of their accurate masses, isotopolog patterns, fragment ions, and retention times (**Table 2**).

To find statistical associations between omics data profiles and colorectal tissue histology, we performed BGA with the full set of metabolites (n=1830) or lipids (n=2365) detected in the tissues. NM, ADN, and CRC samples exhibited markedly different metabolomes and lipidomes (**Figure 1A** and **B**, respectively). The first ordination axis (axis 1), which accounts for the largest proportion of the omics alterations observed during tumorigenesis (quantified as statistical variance), clearly separated the 39 tumors from the 39 matched samples of NM. The ADNs and CRCs were also characterized by distinct metabolomes and lipidomes (axis 2), as were the NM-ADN and NM-CRC samples, as shown in the right-hand graphs in Figure 1.

**Figure 1.**
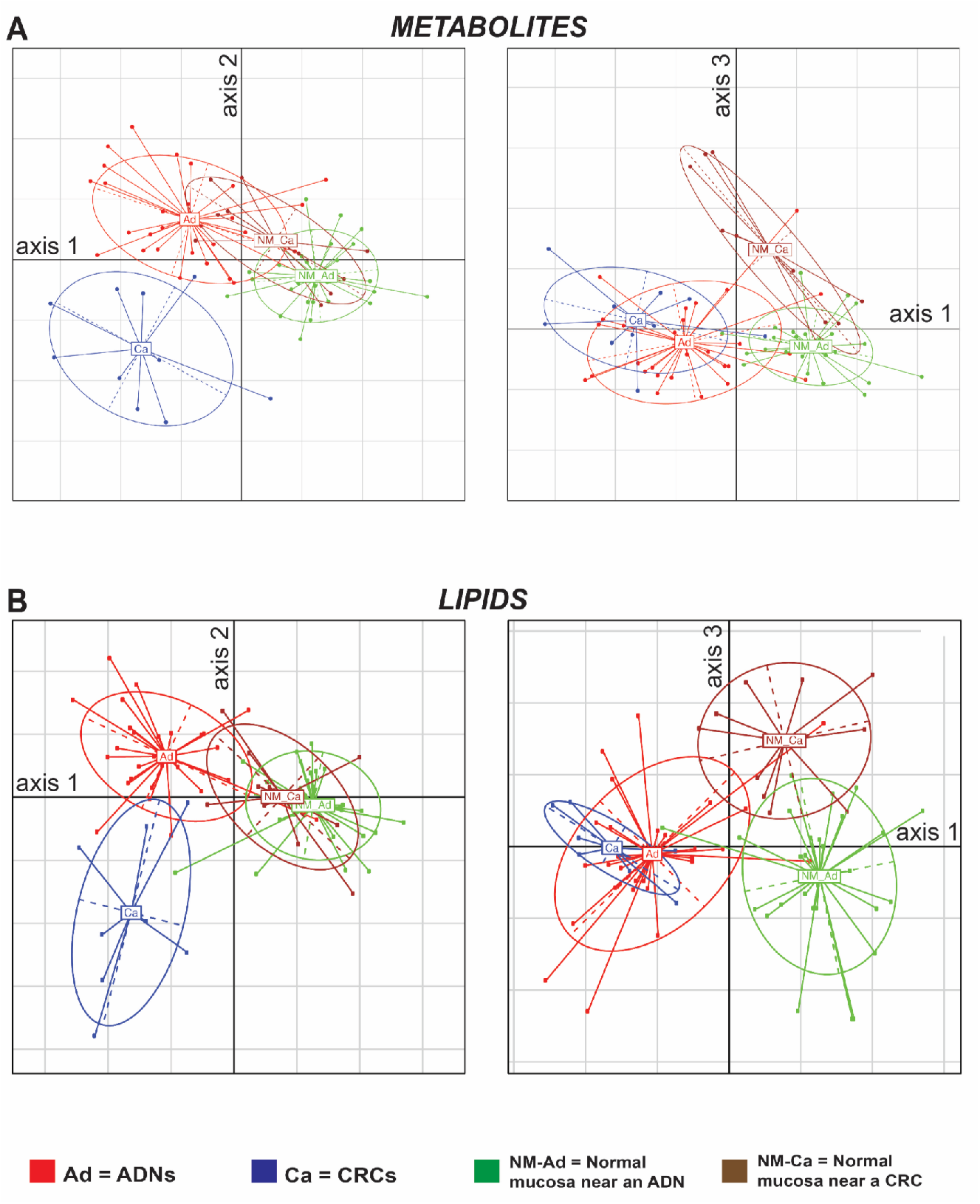
Between-group analysis (BGA) of the 78 colorectal tissue samples based on (A) metabolome and (B) lipidome data. *Left-hand graphs in both panels*: BGA discriminated between normal (NM-ADN and NM-CRC) and tumor (ADN and CRC) samples (axis 1) and between ADNs and CRCs (axis 2) using either data set. Right-hand graphs in both panels show projections along axis 3.

One-way ANOVA was used to determine which detected compounds were significantly more or less abundant in tumor tissues (log2 fold change 1; P value <0.05) than in their matched samples of NM (**Figure 2**). **Supplementary Table 7** (Sheet 1) shows the 261 metabolites that were significantly increased (n=180) or decreased (n=81) in ADNs. Sheet 2 of the same table lists the lipids with higher (n=110) and lower (n=106) levels in the ADNs. Fewer compounds displayed significantly altered levels in CRCs as compared with levels in the NM-CRC samples. As shown in Supplementary Table 7, these included (Sheet 3) 199 metabolites (98 of which had higher levels in CRCs) and (Sheet 4) 148 lipids (53 of which were more abundant in CRCs). Less than one out of five compounds with tumor-associated alterations (11% of the metabolites, 16% of the lipids) were altered in both types of tumor, as shown in the Venn diagrams presented in Figure 2.

**Figure 2.**
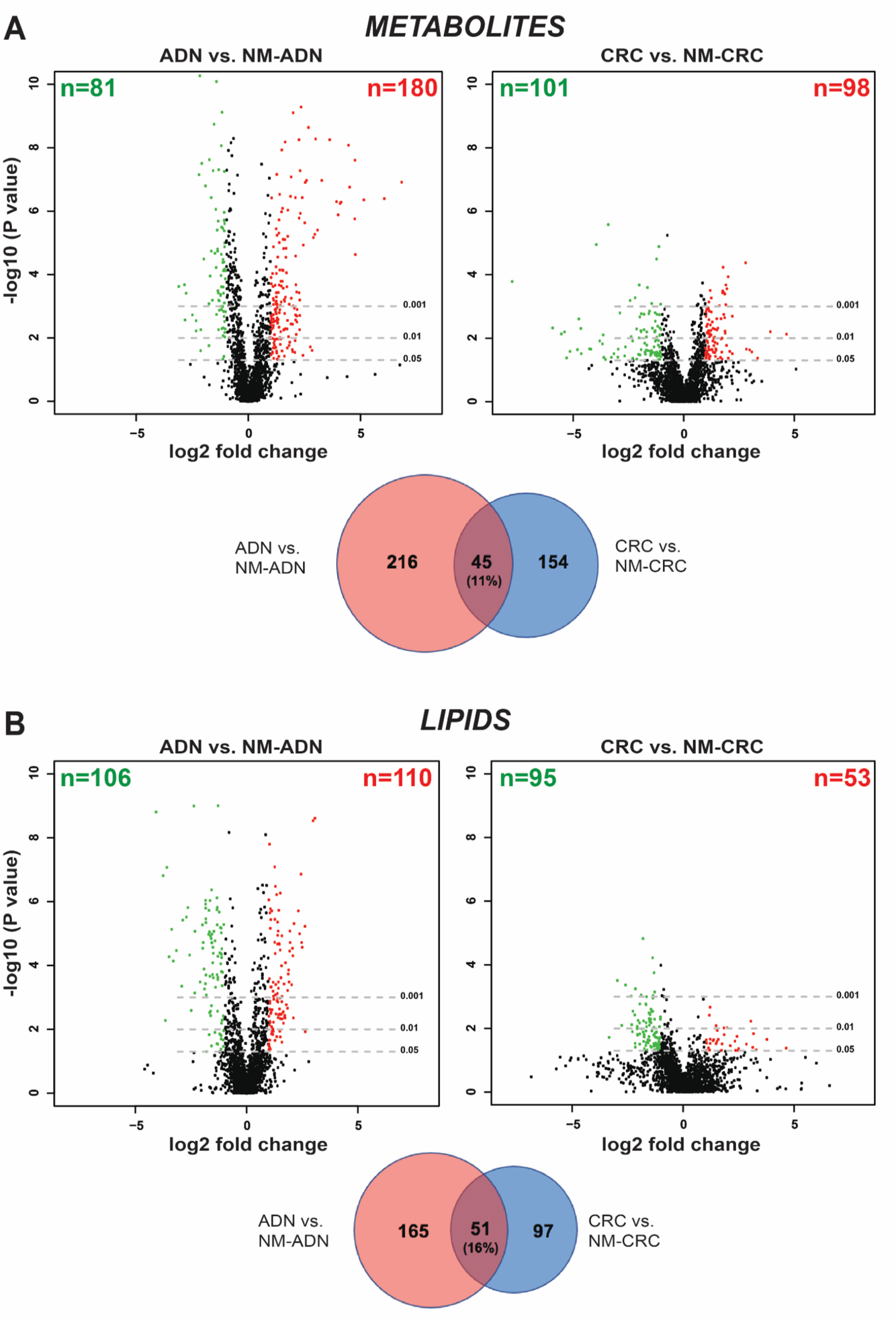
Metabolites and lipids displaying differential abundance in colorectal tumors vs. normal mucosa (NM). Volcano plots were plotted with all the (**A**) metabolites (n=1830) or (**B**) lipids (n=2365) detected in the tissues. Red and green dots represent compounds whose levels in CRCs or ADNs were significantly higher or lower, respectively, than those in tumor-matched samples of NM. Venn diagrams show the number of metabolites (A) and lipids (B) that were differentially abundant in ADNs, in CRCs, and in both tumor types.

Colorectal tumorigenesis is known to involve a progressive accumulation of aberrancies across the transition from NM to ADN and eventually to CRC (see *Introduction*). The relatively small overlap of differentially abundant compounds in ADNs and CRCs and the lower number of significant changes in CRCs than in ADNs were therefore unexpected. However, differences at the metabolome and lipidome levels were also observed between the NM-CRC and NM-ADN samples (Figure 1), as detailed in **Supplementary Figure 1** and **Supplementary Table 8** (Sheets 1 and 2). Compared with NM-CRC, NM-ADN is likely to be more representative of the normal mucosa in a tumor-free colon, since the metabolome and lipidome of non-lesional tissue should be less affected by its proximity to a precancerous rather than a frankly malignant lesion. Indeed, as shown in **Figure 3A** and **Supplementary Figure 2**, when we compared CRCs with NM-ADN, the number of metabolites (n=340) and lipids (n=201) exhibiting cancer-related alterations in abundance (**Supplementary Table 9**, Sheets 1 and 2) clearly exceeded those identified when CRCs were compared with NM-CRCs (n=199 metabolites and n=148 lipids, listed in Supplementary Table 7, Sheets 3 and 4). These findings indicate that many changes occurring in CRCs are statistically undetectable when the comparator is NM-CRC because similar alterations are also present in the apparently normal tissue in the vicinity of a cancer. Invasive tumors like CRCs should also display more changes than their precursor lesions. As shown in **Figure 3B** and **Supplementary Figure 3**, the number of differentially abundant metabolites in CRC—vs. the “more normal” NM-ADN tissues (n=340)—did in fact exceed that found in ADNs (vs NM-ADN) (n=261). The number of differentially abundant lipids in CRCs also increased when the comparator was NM-ADN (n=201, vs. n=148 when compared with NM-CRC [Fig. 2B]) although it still failed to significantly exceed the number identified in ADNs (n=216 [Fig. 3B]) (Supplementary Table 9, Sheets 1 and 2; Supplementary Table 7, Sheets 1 and 2).

**Figure 3.**
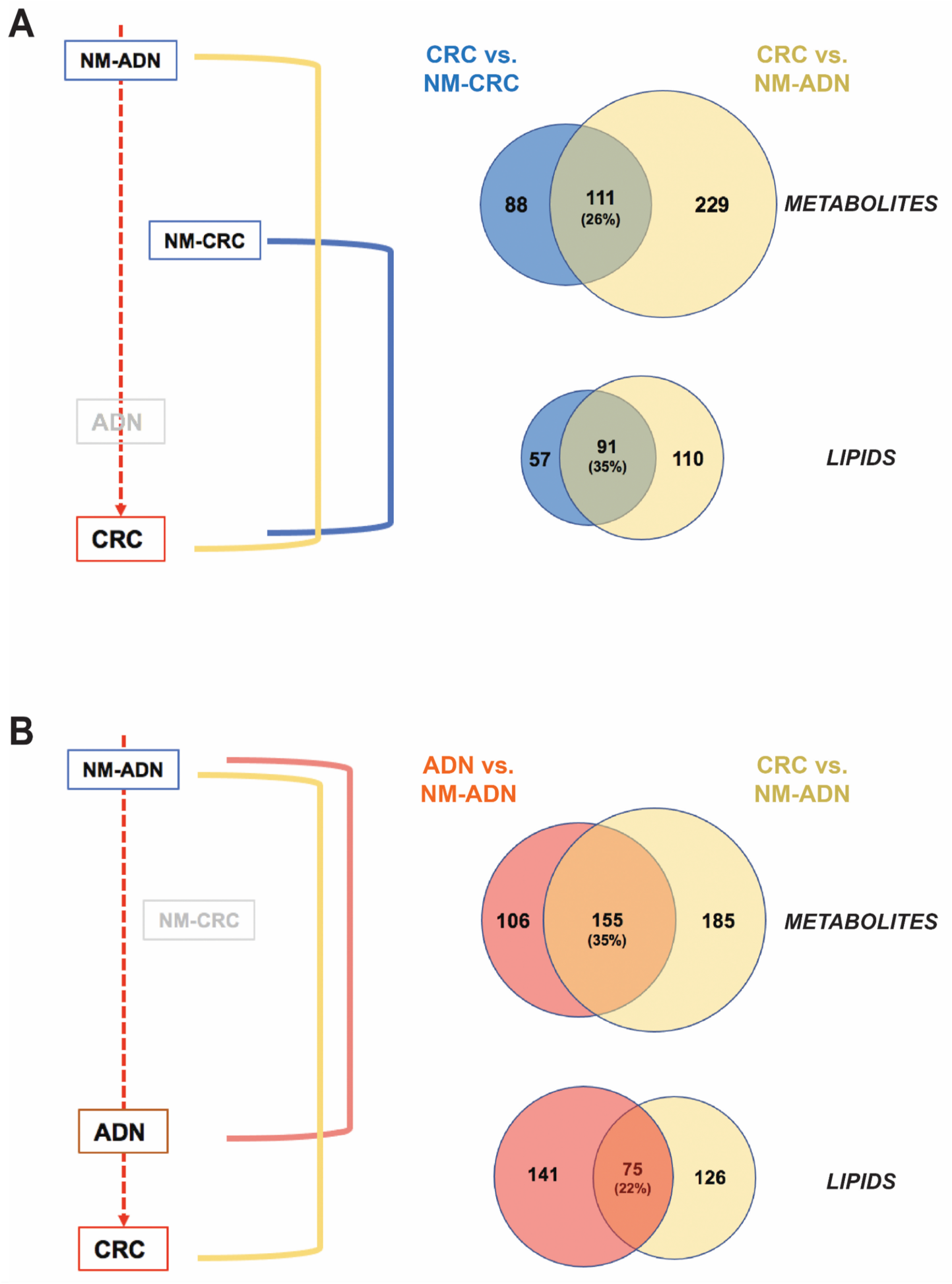
Metabolites and lipids displaying differential abundance in colorectal tumors vs. the two different sets of NM samples. **(A)** As summarized in the diagram on the left, CRCs were compared with NM-CRC samples and with NM-ADN samples. Venn diagrams on the right show the differentially abundant compounds that emerged from each comparison and the overlap between the two sets. (See also the Volcano plots in Supplementary Figure 2). (**B**) Each tumor type was also compared with the NM-AND samples (diagram on the right). As in panel A, Venn diagrams show the differentially abundant compounds that emerged from each comparison and the overlap between the two sets. (See also the Volcano plots in Supplementary Figure 3.)

Nonetheless, use of the NM-ADN samples as the comparator for ADNs as well as CRCs also increased the number of compounds that were identified as altered in both sets of tumors. (Compare set intersections in the Venn diagrams in Figure 3B with those in Figure 2A and B.) This suggests that a considerable number of differentially abundant compounds display *progressively* altered levels across the normal mucosa (as represented in our study by NM-ADN) > adenoma > carcinoma sequence of tumorigenesis. This trend is clearly visible in the heatmaps shown in **Figure 4**: the vast majority of the 340 metabolites and 201 lipids whose abundance in CRCs was significantly higher or lower than that found in the normal mucosa of adenoma patients displayed milder but directionally consistent alterations in the ADNs. As expected on the basis of the BGA data in Figure 1, the NM-CRC samples clustered with the NM-ADN samples (and both were clearly separated from the two tumor types). However, the two NM-CRC and NM-ADN samples also differed from one another in terms of the abundance of several compounds.

**Figure 4.**
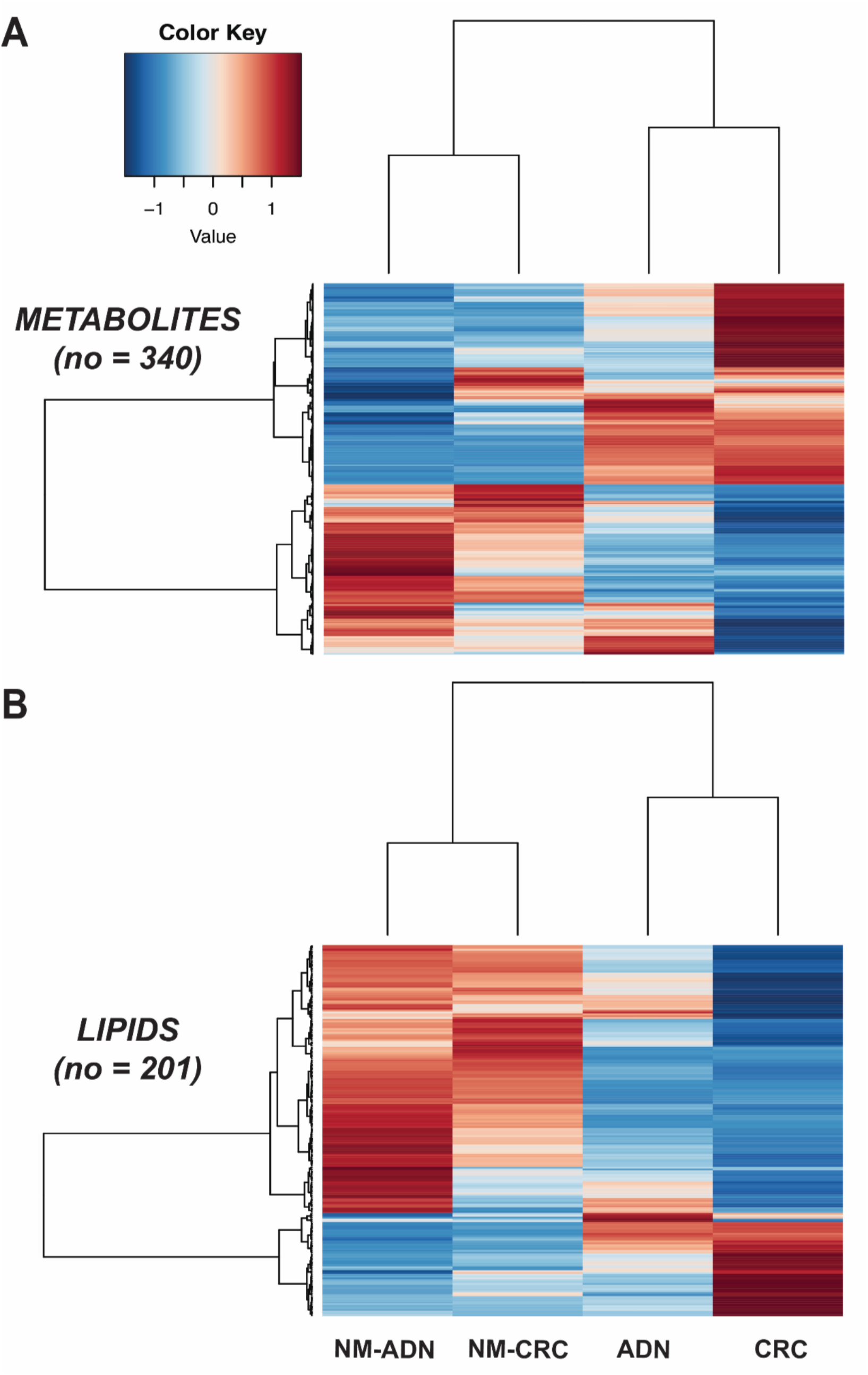
Evolution of metabolite and lipid abundance differentials as colorectal tumorigenesis advances. Heatmaps showing the relative abundance levels of 340 metabolites (**A**) and 201 lipids (**B**) in NM-ADN, NM-CRC, ADNs, and CRCs. Most of the compounds increased progressively or decreased progressively from normal mucosa to ADN and eventually to CRC.

## Discussion

Our untargeted metabolomic and lipidomic studies demonstrate not only that, as expected, the metabolome and lipidome of colorectal tumors differ markedly from those of the normal colorectal mucosa, but also that they evolve with tumor progression. More specifically, most of the metabolites and lipids whose levels are altered in ADNs (as compared with those in the nearby normal-appearing mucosa) undergo further directionally consistent changes in abundance during the subsequent transformation into CRC (Figure 4). Although metabolomic data on colon cancer abounds in the literature, ours are the first to demonstrate the progressive nature of these alterations.

Most of the previously published studies were undertaken to identify diagnostic CRC biomarkers in body fluids (e.g., plasma, serum, stool, or urine), and 42 of these were included in the systematic review and meta-analysis recently conducted by Tian J. et al. ^33^. A smaller, but still impressive, number of studies have been focused on levels of compounds in colorectal tissues ^34–63^. Somewhat surprisingly, there is substantial overlap between the temporal distributions of the two study types in the literature published over the past 15 years. A more rational approach might have begun with the identification of compounds exhibiting substantial abundance differentials in tumor tissues and devoted subsequent studies to the verification of those differences in body fluids. Numerous confounding factors (e.g., age, sex, diet, drugs, co-morbidities) can lead to the false discovery of biomarkers in body fluids ^64–67^. Furthermore, compounds whose levels are increased in specific tumors (as compared with donor-matched normal tissues) may also display increased levels in body fluids, whereas compounds that are less abundant in tumors are poor biofluid biomarker candidates since their decreased tumor levels are very unlikely to be reflected in the body fluids. To our knowledge, only one attempt has been made thus far to identify metabolites whose levels are altered in both the cancer tissue and body-fluid compartments (specifically, in plasma) ^68^. The authors identified 8 “putative biomarkers,” but some of these displayed directionally opposite changes in abundance in plasma and CRCs (including L-tryptophan, L-tyrosine, and chenodeoxycholic acid: compare Supplementary Table S4 and S6 of ^68^). Of course, cancer-mediated alterations in the abundance of a compounds in body fluids can be identified regardless of whether or not there are metabolomic changes in the cancer tissue itself: such changes can easily result from the systemic consequences of an invasive tumor (e.g., responses to the cancer by the patient’s immune system or other organs). And some of these changes might even display directional inconsistency in cancer tissue and body fluid compartments. The fact that such alterations are more likely to occur during the advanced stage of CRC would certainly limit their value for the early diagnosis of cancer and virtually exclude their use for the detection of colorectal adenomas and other non-invasive, precancerous tumors of the gut.

The studies cited above were all conducted in CRCs, although precancerous lesions were also occasionally included in the analyses. In the study by Tessem et al., HR-MAS NMR was used to analyze 4 adenomas, 28 CRCs, and matched samples of normal mucosa. In principal component analysis, the precancerous lesions could be distinguished from the CRCs, mainly by their higher levels of phosphocholine. Ong et al. analyzed 26 “polyps” from patients who also had CRC, but no clinical or pathological information was provided on the precancerous lesions. (Curiously, this 2010 publication continues to appear as an article *in press* in Molecular and Cellular Proteomics, and for this reason, it is not included in our *References* section). Shureiqi et al. ^40^ used LC-MS/MS to assess levels of six compounds produced during lipoxygenase metabolism in 36 colorectal “polyps” and 40 CRCs and found 13-hydroxyoctadecadienoic acid levels to be similarly decreased in both types of lesion, as compared with normal mucosa. In the study by Gao et al., ^55^ 10 adenomas were included in a targeted analysis of amino acid profiles using UPLC-MS. These precancerous lesions were compared with CRCs, but not with matched samples of normal mucosa. However, the adenomas and CRCs were found to differ significantly in terms of their content of nine amino acids. Satoh et al. ^69^ included only 5 adenomas in their analysis of 275 colorectal tumors using CE-TOF-MS, and the metabolome of the few precancerous lesions could not be distinguished from that of the CRCs. Finally, HR-MAS NMR was used by Kinross et al. ^58^ to analyze 18 surgically resected colorectal tumors from 18 patients. Three of the lesions proved to be large adenomas (which frequently display high grade dysplasia, although this feature was not specified in the report), and the metabolome of these lesions was apparently not significantly different from that of the 15 CRCs.

Compared with the metabolome findings published thus far on colorectal adenomas, ours is the first study that systematically compared the contents of more than 4000 metabolites and lipids in adenomas, CRCs, and tumor-matched sample of normal mucosa. To this end, we used nano-flow LC-MS (inner column diameter of 200 µm), ran each sample in both HILIC and RP modes, and analyzed the acquired LC-MS/MS data with Progenesis QI (see *Methods*). Importantly, the tissues we studied were collected during endoscopy, and within a few seconds of excision, each sample was frozen in liquid nitrogen to halt enzymatic reactions. This contrasts with the methods used in all previously published studies, where tissue samples were taken from surgical specimens exposed to variable periods of intraoperative ischemia, which can alter metabolite levels ^70^, and variable intervals between resection and sample freezing, which can affect tissue integrity.

In our study, the metabolome and lipidome of normal-appearing colorectal mucosa samples taken 2-5 cm from a CRC differed, at both the metabolomic and lipidomic levels, from normal mucosa samples collected near an adenoma. This finding might be a reflection of the field cancerization phenomenon that has been described in CRC. In previous studies, the metabolome of normal-appearing colorectal mucosa from patients with CRC has been found to differ with both the type and stage of the cancer ^41,46,71^, so it is not surprising that normal mucosa in the vicinity of a precancerous colorectal lesion would differ from that taken near one that has already undergone transformation. The absence in our study of colorectal tissues from donors with lesion-free colons limited our ability to explore this hypothesis. However, the putative role of a field effect is consistent not only with previous reports in the literature but also with the findings that emerged from our comparison of CRCs with NM-ADN.

The main limitation of our study is the lack of definitive annotations for most of the compounds detected in the tissues. As a result, we were unable to perform metabolic pathway analysis. However, the raw data we have deposited in the *Metabolights* repository can springboard further analyses aimed at the definitive identification of metabolites and lipids typically abundant in colorectal tumors. Such compounds will be promising candidates for the development of diagnostic tumor biomarkers, and they can also be expected to shed valuable light on the metabolic landscape in adenomas and CRCs. Integrated with other types of omics data on colorectal tissues collected (by our groups and others) over the last decade or so, this knowledge will hopefully bring us closer to obtaining a comprehensive picture of the metabolic abnormalities in these tumors, a goal that continues to elude us today in spite of the numerous studies that have already been published in the field ^33,72^. Achieving this goal will also be a major step forward on the road toward the development of more effective therapeutic interventions against CRC.

## List of abbreviations

ADN: conventional colorectal adenoma
BGA: Between Group Analysis
CRC: colorectal cancer
ESI: electro-spray ionization
HILIC: Hydrophilic Interaction Liquid Chromatography
LC-MS: Liquid Chromatography - Mass Spectrometry
MS: Mass Spectrometry
NM: normal mucosa
NM-ADN: NM collected near a colorectal adenoma
NM-CRC: NM collected near a CRC
RP: Reversed Phase

## Ethical approval and consent to participate

This study was approved by the Zurich Cantonal Ethics Committee (KEK-2011-0275) and informed consent was obtained from all individual patients included in the study.

## Consent for publication

Not applicable

## Authors’ contributions

EL and GM designed the experiments, analyzed the data, and wrote the manuscript. CM collected tissue samples and clinical data.

## Acknowledgements

We thank David Fischer for technical assistance in processing samples and producing the raw data that were utilized in this study. We also thank Marian Everett Kent for editing the manuscript.

## Supplementary Material

**Supplementary Figure 1. Metabolites and lipids displaying differential abundance in normal mucosa near an adenoma (NM-ADN) versus that near a CRC (NM-CRC)**. Volcano plots show (**A**) metabolites and (**B**) lipids whose levels in NM-CRC were significantly higher (red dots) or lower (green dots) than those in NM-ADN.

**Supplementary Figure 2. Metabolites and lipids displaying differential abundance in CRCs as compared with NM-CRC samples or with NM-ADN samples**. Volcano plots showing the numbers of metabolites (**A**) and lipids (**B**) whose levels in CRCs were significantly increased (red dots) or decreased (green dots) in the two comparisons. Venn diagrams show the overlap between the two sets of differentially abundant compounds.

**Supplementary Figure 3. Metabolites and lipids displaying differential abundance in ADNs vs. NM-ADN and in CRCs vs. NM-ADN**. Volcano plots showing the number of significantly more (red dots) or less (green dots) abundant metabolites (**A**) or lipids (**B**) in ADNs and CRCs. Venn diagrams show the overlap between the two sets of differentially abundant compounds.

**Supplementary Table 1**. Composition of the internal standards mixture added to tissue samples prior to LC-MS.

**Supplementary Table 2**. Details of the two LC-MS methods used in the study.

**Supplementary Table 3**. Core Metabolites. Full list of the 190 well characterized, and reliably detectable and quantifiable metabolites.

**Supplementary Table 4**. Sample descriptors.

**Supplementary Table 5**. Metabolite-intensity data and best matching annotations.

**Supplementary Table 6**. Lipid-intensity data and best matching annotations.

**Supplementary Table 7** (4 Sheets). Differentially abundant compounds in tumors (in comparison with their matched samples of normal mucosa).

**Supplementary Table 8**. Sheet 1, metabolites displaying differential abundance in NM-CRC samples vs. NM-ADN samples. Sheet 2, lipids displaying differential abundance in NM-CRC samples vs. NM-ADN samples.

**Supplementary Table 9**. (2 Sheets). Differentially abundant compounds in CRCs (in comparison with normal mucosa samples of adenoma patients).

## References

1. Arndt V, Feller A, Hauri D, Heusser R, Junker C, Kuehni C, Lorez M, Pfeiffer V, Roy E, Schindler M. Swiss Cancer Report 2015. Neuchatel, Switzerland: Federal Statistical Office; 2016;

2. Winawer SJ. Natural history of colorectal cancer. Am J Medicine 1999; 106:3–6.

3. Lash RH, Genta RM, Schuler CM. Sessile serrated adenomas: prevalence of dysplasia and carcinoma in 2139 patients. J Clin Pathol 2010; 63:681.

4. Bettington M, Walker N, Rosty C, Brown I, Clouston A, McKeone D, Pearson S-A, Leggett B, Whitehall V. Clinicopathological and molecular features of sessile serrated adenomas with dysplasia or carcinoma. Gut 2017; 66:97.

5. Meester RGS, Herk MMAGC van, Lansdorp-Vogelaar I, Ladabaum U. Prevalence and Clinical Features of Sessile Serrated Polyps: A Systematic Review. Gastroenterology 2020; 159:105-118.e25.

6. Carr PR, Weigl K, Edelmann D, Jansen L, Chang-Claude J, Brenner H, Hoffmeister M. Estimation of Absolute Risk of Colorectal Cancer Based on Healthy Lifestyle, Genetic Risk, and Colonoscopy Status in a Population-Based Study. Gastroenterology 2020; 159:129-138.e9.

7. Mendelsohn RB, Winawer SJ, Ahnen DJ. Incidence of Colorectal Cancer Matters. Gastroenterology 2019; 158:1191–5.

8. Spring KJ, Zhao ZZ, Karamatic R, Walsh MD, Whitehall VLJ, Pike T, Simms LA, Young J, James M, Montgomery GW, et al. High Prevalence of Sessile Serrated Adenomas With BRAF Mutations: A Prospective Study of Patients Undergoing Colonoscopy. Gastroenterology 2006; 131:1400–7.

9. Snover DC. Diagnostic and reporting issues of preneoplastic polyps of the large intestine with early carcinoma. Ann Diagn Pathol 2018; 39:1–14.

10. Brenner H, Hoffmeister M, Stegmaier C, Brenner G, Altenhofen L, Haug U. Risk of progression of advanced adenomas to colorectal cancer by age and sex: estimates based on 840 149 screening colonoscopies. Gut 2007; 56:1585–9.

11. Hassan C, Quintero E, Dumonceau J-M, Regula J, Brandão C, Chaussade S, Dekker E, Dinis-Ribeiro M, Ferlitsch M, Gimeno-García A, et al. Post-polypectomy colonoscopy surveillance: European Society of Gastrointestinal Endoscopy (ESGE) Guideline. Endoscopy 2013; 45:842–64.

12. Kaltenbach T, Anderson JC, Burke CA, Dominitz JA, Gupta S, Lieberman D, Robertson DJ, Shaukat A, Syngal S, Rex DK. Endoscopic Removal of Colorectal Lesions—Recommendations by the US Multi-Society Task Force on Colorectal Cancer. Gastroenterology 2020; 158:1095–129.

13. Pai RK, Bettington M, Srivastava A, Rosty C. An update on the morphology and molecular pathology of serrated colorectal polyps and associated carcinomas. Modern Pathol 2019; 32:1390–415.

14. Vogelstein B, Fearon ER, Hamilton SR, Kern SE, Preisinger AC, Leppert M, Nakamura Y, White R, Smits AM, Bos JL. Genetic Alterations during Colorectal-Tumor Development. New Engl J Medicine 1988; 319:525–32.

15. Wood LD, Parsons DW, Jones S, Lin J, Sjöblom T, Leary RJ, Shen D, Boca SM, Barber T, Ptak J, et al. The Genomic Landscapes of Human Breast and Colorectal Cancers. Science 2007; 318:1108–13.

16. Muzny DM, Bainbridge MN, Chang K, Dinh HH, Drummond JA, Fowler G, Kovar CL, Lewis LR, Morgan MB, Newsham IF, et al. Comprehensive molecular characterization of human colon and rectal cancer. Nature 2012; 487:330–7.

17. Nikolaev SI, Sotiriou SK, Pateras IS, Santoni F, Sougioultzis S, Edgren H, Almusa H, Robyr D, Guipponi M, Saarela J, et al. A Single-Nucleotide Substitution Mutator Phenotype Revealed by Exome Sequencing of Human Colon Adenomas. Cancer Res 2012; 72:6279–89.

18. Kleeman SO, Koelzer VH, Jones HJ, Vazquez EG, Davis H, East JE, Arnold R, Koppens MA, Blake A, Domingo E, et al. Exploiting differential Wnt target gene expression to generate a molecular biomarker for colorectal cancer stratification. Gut 2020; 69:1092– 103.

19. Clevers H. Wnt/β-Catenin Signaling in Development and Disease. Cell 2006; 127:469–80.

20. Sabates-Bellver J, Flier LGV der, Palo M de, Cattaneo E, Maake C, Rehrauer H, Laczko E, Kurowski MA, Bujnicki JM, Menigatti M, et al. Transcriptome Profile of Human Colorectal Adenomas. Mol Cancer Res 2007; 5:1263–75.

21. Van der Flier LG, Sabates–Bellver J, Oving I, Haegebarth A, Palo MD, Anti M, Gijn MEV, Suijkerbuijk S, Wetering MV de, Marra G, et al. The Intestinal Wnt/TCF Signature. Gastroenterology 2007; 132:628–32.

22. Cattaneo E, Laczko E, Buffoli F, Zorzi F, Bianco MA, Menigatti M, Bartosova Z, Haider R, Helmchen B, Sabates-Bellver J, et al. Preinvasive colorectal lesion transcriptomes correlate with endoscopic morphology (polypoid vs. nonpolypoid). Embo Mol Med 2011; 3:334–47.

23. Vonlanthen J, Okoniewski MJ, Menigatti M, Cattaneo E, Pellegrini-Ochsner D, Haider R, Jiricny J, Staiano T, Buffoli F, Marra G. A comprehensive look at transcription factor gene expression changes in colorectal adenomas. Bmc Cancer 2014; 14:46.

24. di Pietro M, Sabates-Bellver J, Menigatti M, Bannwart F, Schnider A, Russell A, Truninger K, Jiricny J, Marra G. Defective DNA Mismatch Repair Determines a Characteristic Transcriptional Profile in Proximal Colon Cancers. Gastroenterology 2005; 129:1047–59.

25. Parker HR, Orjuela S, Oliveira AM, Cereatti F, Sauter M, Heinrich H, Tanzi G, Weber A, Komminoth P, Vavricka S, et al. The proto CpG island methylator phenotype of sessile serrated adenomas/polyps. Epigenetics 2018; 13:1088–105.

26. Noreen F, Küng T, Tornillo L, Parker H, Silva M, Weis S, Marra G, Rad R, Truninger K, Schär P. DNA methylation instability by BRAF-mediated TET silencing and lifestyle-exposure divides colon cancer pathways. Clin Epigenetics 2019; 11:196.

27. Uzozie A, Nanni P, Staiano T, Grossmann J, Barkow-Oesterreicher S, Shay JW, Tiwari A, Buffoli F, Laczko E, Marra G. Sorbitol Dehydrogenase Overexpression and Other Aspects of Dysregulated Protein Expression in Human Precancerous Colorectal Neoplasms: A Quantitative Proteomics Study. Mol Cell Proteomics 2014; 13:1198–218.

28. Uzozie AC, Selevsek N, Wahlander A, Nanni P, Grossmann J, Weber A, Buffoli F, Marra G. Targeted Proteomics for Multiplexed Verification of Markers of Colorectal Tumorigenesis. Mol Cell Proteomics 2017; 16:407–27.

29. World Health Organization. Classification of tumours of the digestive tract. Lyon: IARC Press; 2019.

30. Plumb RS, Johnson KA, Rainville P, Smith BW, Wilson ID, Castro-Perez JM, Nicholson JK. UPLC/MSE; a new approach for generating molecular fragment information for biomarker structure elucidation. Rapid Commun Mass Sp 2006; 20:1989–94.

31. Culhane AC, Perrière G, Considine EC, Cotter TG, Higgins DG. Between-group analysis of microarray data. Bioinformatics 2002; 18:1600–8.

32. Culhane AC, Thioulouse J, Perrière G, Higgins DG. MADE4: an R package for multivariate analysis of gene expression data. Bioinformatics 2005; 21:2789–90.

33. Tian J, Xue W, Yin H, Zhang N, Zhou J, Long Z, Wu C, Liang Z, Xie K, Li S, et al. Differential Metabolic Alterations and Biomarkers Between Gastric Cancer and Colorectal Cancer: A Systematic Review and Meta-Analysis. Oncotargets Ther 2020; 13:6093–108.

34. Denkert C, Budczies J, Weichert W, Wohlgemuth G, Scholz M, Kind T, Niesporek S, Noske A, Buckendahl A, Dietel M, et al. Metabolite profiling of human colon carcinoma – deregulation of TCA cycle and amino acid turnover. Mol Cancer 2008; 7:72.

35. Piotto M, Moussallieh F-M, Dillmann B, Imperiale A, Neuville A, Brigand C, Bellocq J-P, Elbayed K, Namer IJ. Metabolic characterization of primary human colorectal cancers using high resolution magic angle spinning 1H magnetic resonance spectroscopy. Metabolomics 2008; 5:292–301.

36. Hirayama A, Kami K, Sugimoto M, Sugawara M, Toki N, Onozuka H, Kinoshita T, Saito N, Ochiai A, Tomita M, et al. Quantitative Metabolome Profiling of Colon and Stomach Cancer Microenvironment by Capillary Electrophoresis Time-of-Flight Mass Spectrometry. Cancer Res 2009; 69:4918–25.

37. Righi V, Durante C, Cocchi M, Calabrese C, Febo GD, Lecce F, Pisi A, Tugnoli V, Mucci A, Schenetti L. Discrimination of Healthy and Neoplastic Human Colon Tissues by ex Vivo HR-MAS NMR Spectroscopy and Chemometric Analyses. J Proteome Res 2009; 8:1859–69.

38. Chan ECY, Koh PK, Mal M, Cheah PY, Eu KW, Backshall A, Cavill R, Nicholson JK, Keun HC. Metabolic Profiling of Human Colorectal Cancer Using High-Resolution Magic Angle Spinning Nuclear Magnetic Resonance (HR-MAS NMR) Spectroscopy and Gas Chromatography Mass Spectrometry (GC/MS). J Proteome Res 2009; 8:352–61.

39. Chae Y-K, Kang W-Y, Kim S-H, Joo J-E, Han J-K, Hong B-W. Combining Information of Common Metabolites Reveals Global Differences between Colorectal Cancerous and Normal Tissues. B Korean Chem Soc 2010; 31:379–83.

40. Shureiqi I, Chen D, Day RS, Zuo X, Hochman FL, Ross WA, Cole RA, Moy O, Morris JS, Xiao L, et al. Profiling Lipoxygenase Metabolism in Specific Steps of Colorectal Tumorigenesis. Cancer Prev Res 2010; 3:829–38.

41. Tessem M-B, Selnæs KM, Sjursen W, Tranø G, Giskeødegård GF, Bathen TF, Gribbestad IS, Hofsli E. Discrimination of Patients with Microsatellite Instability Colon Cancer using 1 H HR MAS MR Spectroscopy and Chemometric Analysis. J Proteome Res 2010; 9:3664–70.

42. Mal M, Koh PK, Cheah PY, Chan ECY. Metabotyping of human colorectal cancer using two-dimensional gas chromatography mass spectrometry. Anal Bioanal Chem 2012; 403:483–93.

43. Kim S, Lee S, Maeng YH, Chang WY, Hyun JW, Kim S. Study of Metabolic Profiling Changes in Colorectal Cancer Tissues Using 1D H-1 HR-MAS NMR Spectroscopy. B Korean Chem Soc 2013; 34:1467–72.

44. Holst S, Stavenhagen K, Balog CIA, Koeleman CAM, McDonnell LM, Mayboroda OA, Verhoeven A, Mesker WE, Tollenaar Raem, Deelder AM, et al. Investigations on Aberrant Glycosylation of Glycosphingolipids in Colorectal Cancer Tissues Using Liquid Chromatography and Matrix-Assisted Laser Desorption Time-of-Flight Mass Spectrometry (MALDI-TOF-MS). Mol Cell Proteomics 2013; 12:3081–93.

45. Wang H, Wang L, Zhang H, Deng P, Chen J, Zhou B, Hu J, Zou J, Lu W, Xiang P, et al. 1H NMR-based metabolic profiling of human rectal cancer tissue. Mol Cancer 2013; 12:121.

46. Jiménez B, Mirnezami R, Kinross J, Cloarec O, Keun HC, Holmes E, Goldin RD, Ziprin P, Darzi A, Nicholson JK. 1 H HR-MAS NMR Spectroscopy of Tumor-Induced Local Metabolic “Field-Effects” Enables Colorectal Cancer Staging and Prognostication. J Proteome Res 2013; 12:959–68.

47. Manna SK, Tanaka N, Krausz KW, Haznadar M, Xue X, Matsubara T, Bowman ED, Fearon ER, Harris CC, Shah YM, et al. Biomarkers of Coordinate Metabolic Reprogramming in Colorectal Tumors in Mice and Humans. Gastroenterology 2014; 146:1313–24.

48. Phua LC, Chue XP, Koh PK, Cheah PY, Ho HK, Chan ECY. Non-invasive fecal metabonomic detection of colorectal cancer. Cancer Biol Ther 2014; 15:389– 97.

49. Qiu Y, Cai G, Zhou B, Li D, Zhao A, Xie G, Li H, Cai S, Xie D, Huang C, et al. A Distinct Metabolic Signature of Human Colorectal Cancer with Prognostic Potential. Clin Cancer Res 2014; 20:2136–46.

50. Mirnezami R, Jiménez B, Li JV, Kinross JM, Veselkov K, Goldin RD, Holmes E, Nicholson JK, Darzi A. Rapid Diagnosis and Staging of Colorectal Cancer via High-Resolution Magic Angle Spinning Nuclear Magnetic Resonance (HR-MAS NMR) Spectroscopy of Intact Tissue Biopsies. Ann Surg 2014; 259:1138–49.

51. Williams MD, Zhang X, Park J-J, Siems WF, Gang DR, Resar LMS, Reeves R, Hill HH. Characterizing metabolic changes in human colorectal cancer. Anal Bioanal Chem 2015; 407:4581–95.

52. Johnson CH, Dejea CM, Edler D, Hoang LT, Santidrian AF, Felding BH, Ivanisevic J, Cho K, Wick EC, Hechenbleikner EM, et al. Metabolism Links Bacterial Biofilms and Colon Carcinogenesis. Cell Metab 2015; 21:891–7.

53. Zhang H, Qiao L, Li X, Wan Y, Yang L, Wang H. Tissue metabolic profiling of lymph node metastasis of colorectal cancer assessed by 1H NMR. Oncol Rep 2016; 36:3436–48.

54. Tian Y, Xu T, Huang J, Zhang L, Xu S, Xiong B, Wang Y, Tang H. Tissue Metabonomic Phenotyping for Diagnosis and Prognosis of Human Colorectal Cancer. Sci Rep. 2016; 6:20790.

55. Gao P, Zhou C, Zhao L, Zhang G, Zhang Y. Tissue amino acid profile could be used to differentiate advanced adenoma from colorectal cancer. J Pharm Biomed Anal 2016; 118:349–55.

56. Ning W, Li H, Meng F, Cheng J, Song X, Zhang G, Wang W, Wu S, Fang J, Ma K, et al. Identification of differential metabolic characteristics between tumor and normal tissue from colorectal cancer patients by gas chromatography–mass spectrometry. Biomed Chromatogr 2017; 31:e3999.

57. Mika A, Kobiela J, Czumaj A, Chmielewski M, Stepnowski P, Sledzinski T. Hyper-Elongation in Colorectal Cancer Tissue – Cerotic Acid is a Potential Novel Serum Metabolic Marker of Colorectal Malignancies. Cell Physiol Biochem 2017; 41:722–30.

58. Kinross J, Mirnezami R, Alexander J, Brown R, Scott A, Galea D, Veselkov K, Goldin R, Darzi A, Nicholson J, et al. A prospective analysis of mucosal microbiome-metabonome interactions in colorectal cancer using a combined MAS 1HNMR and metataxonomic strategy. Sci Rep 2017; 7:8979.

59. Lin Y, Ma C, Bezabeh T, Wang Z, Liang J, Huang Y, Zhao J, Liu X, Ye W, Tang W, et al. 1H NMR-based metabolomics reveal overlapping discriminatory metabolites and metabolic pathway disturbances between colorectal tumor tissues and fecal samples. Int J Cancer 2019; 145:1679–89.

60. Wang Y, Hinz S, Uckermann O, Hönscheid P, Schönfels W von, Burmeister G, Hendricks A, Ackerman JM, Baretton GB, Hampe J, et al. Shotgun lipidomics-based characterization of the landscape of lipid metabolism in colorectal cancer. Biochim Biophys Acta - Mol Cell Biol Lipids 2019; 1865:158579.

61. Long Z, Zhou J, Xie K, Wu Z, Yin H, Daria V, Tian J, Zhang N, Li L, Zhao Y, et al. Metabolomic Markers of Colorectal Tumor With Different Clinicopathological Features. Frontiers Oncol 2020; 10:981.

62. Cai Y, Rattray NJW, Zhang Q, Mironova V, Santos-Neto A, Muca E, Vollmar AKR, Hsu K-S, Rattray Z, Cross JR, et al. Tumor Tissue-Specific Biomarkers of Colorectal Cancer by Anatomic Location and Stage. Metabolites 2020; 10:257.

63. Cai Y, Rattray NJW, Zhang Q, Mironova V, Santos-Neto A, Hsu K-S, Rattray Z, Cross JR, Zhang Y, Paty PB, et al. Sex Differences in Colon Cancer Metabolism Reveal A Novel Subphenotype. Sci Rep 2020; 10:4905.

64. Li Y, Li M, Jia W, Ni Y, Chen T. MCEE: a data preprocessing approach for metabolic confounding effect elimination. Anal Bioanal Chem 2018; 410:2689–99.

65. Cross AJ, Moore SC, Boca S, Huang W, Xiong X, Stolzenberg-Solomon R, Sinha R, Sampson JN. A prospective study of serum metabolites and colorectal cancer risk. Cancer 2014; 120:3049–57.

66. Moore SC, Matthews CE, Sampson JN, Stolzenberg-Solomon RZ, Zheng W, Cai Q, Tan YT, Chow W-H, Ji B-T, Liu DK, et al. Human metabolic correlates of body mass index. Metabolomics 2014; 10:259–69.

67. Slupsky CM, Rankin KN, Wagner J, Fu H, Chang D, Weljie AM, Saude EJ, Lix B, Adamko DJ, Shah S, et al. Investigations of the Effects of Gender, Diurnal Variation, and Age in Human Urinary Metabolomic Profiles. Anal Chem 2007; 79:6995–7004.

68. Wang Z, Cui B, Zhang F, Yang Y, Shen X, Li Z, Zhao W, Zhang Y, Deng K, Rong Z, et al. Development of a Correlative Strategy To Discover Colorectal Tumor Tissue Derived Metabolite Biomarkers in Plasma Using Untargeted Metabolomics. Anal Chem 2018; 91:2401–8.

69. Satoh K, Yachida S, Sugimoto M, Oshima M, Nakagawa T, Akamoto S, Tabata S, Saitoh K, Kato K, Sato S, et al. Global metabolic reprogramming of colorectal cancer occurs at adenoma stage and is induced by MYC. Proc Natl Acad Sci 2017; 114:E7697–706.

70. Cacciatore S, Hu X, Viertler C, Kap M, Bernhardt GA, Mischinger H-J, Riegman P, Zatloukal K, Luchinat C, Turano P. Effects of Intra-and Post-Operative Ischemia on the Metabolic Profile of Clinical Liver Tissue Specimens Monitored by NMR. J Proteome Res 2013; 12:5723–9.

71. Lean CL, Newland RC, Ende DA, Bokey EL, Smith ICP, Mountford CE. Assessment of human colorectal biopsies by 1H MRS: Correlation with histopathology. Magnet Reson Med 1993; 30:525–33.

72. Williams MD, Reeves R, Resar LS, Hill HH. Metabolomics of colorectal cancer: past and current analytical platforms. Anal Bioanal Chem 2013; 405:5013–30.

