## Supplementary figures and images for "The metabolome and lipidome of colorectal adenomas and cancers"

### Supplementary Figure 1

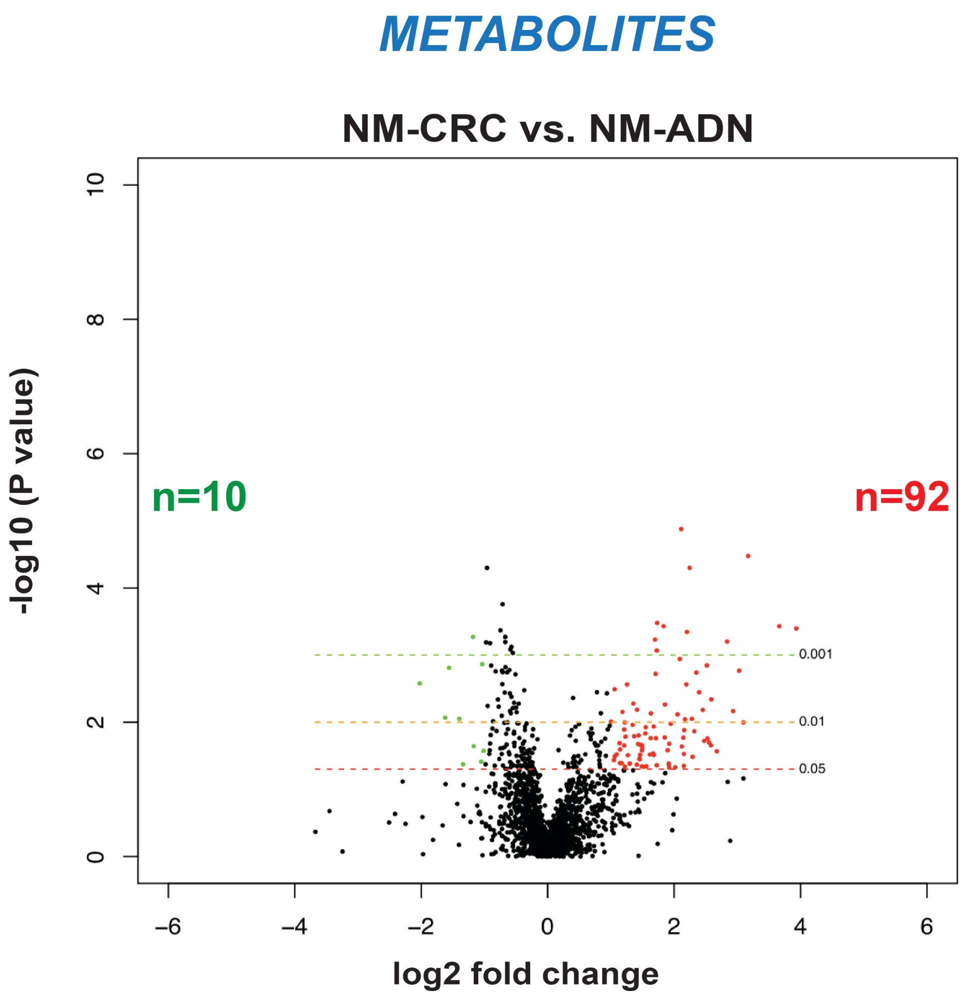

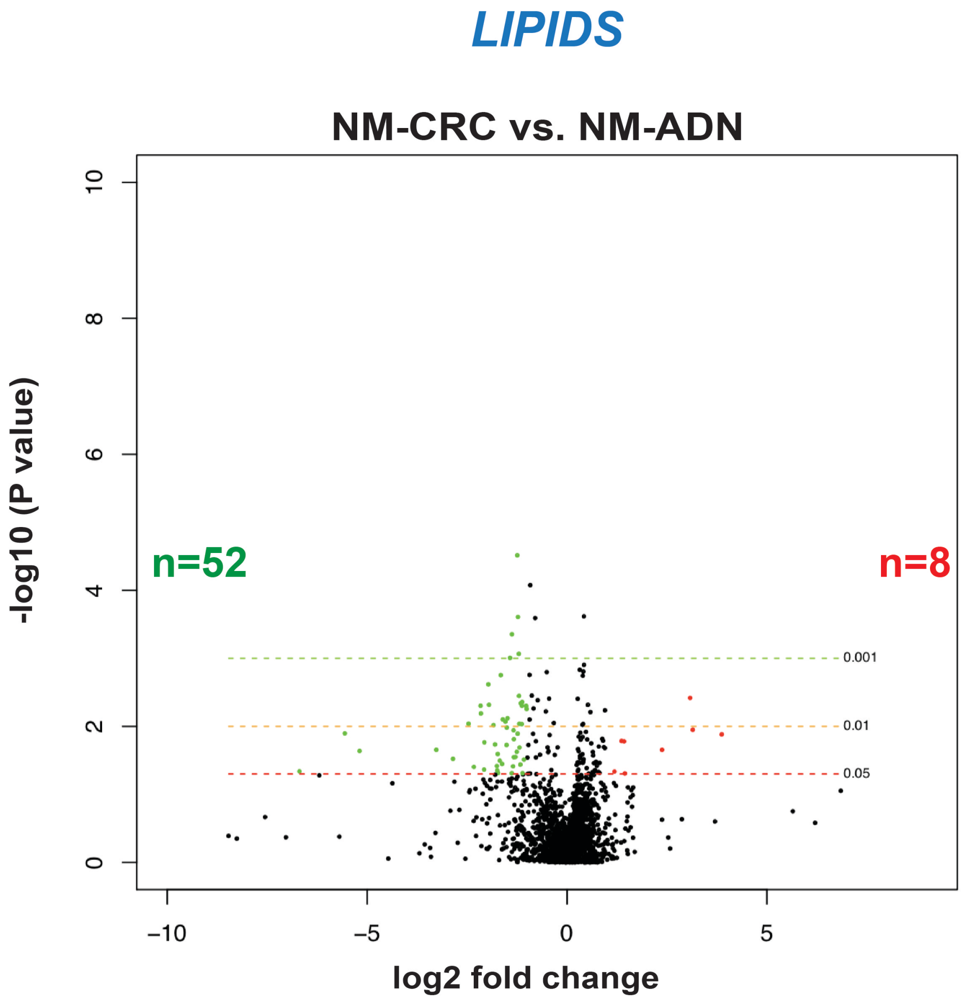

### Supplementary Figure 2

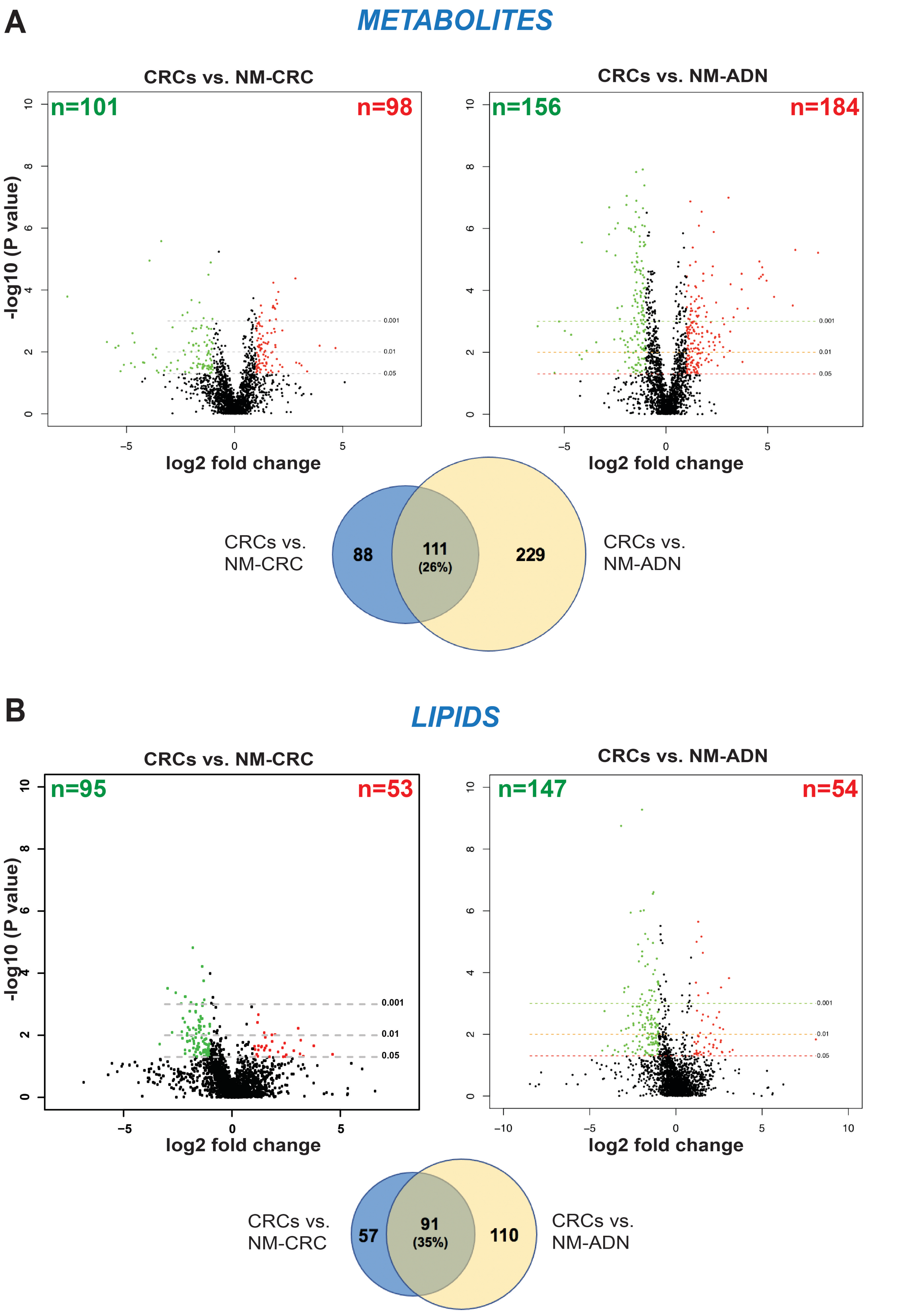

### Supplementary Figure 3

Supplementary Figure 3, Laczko E. et al.

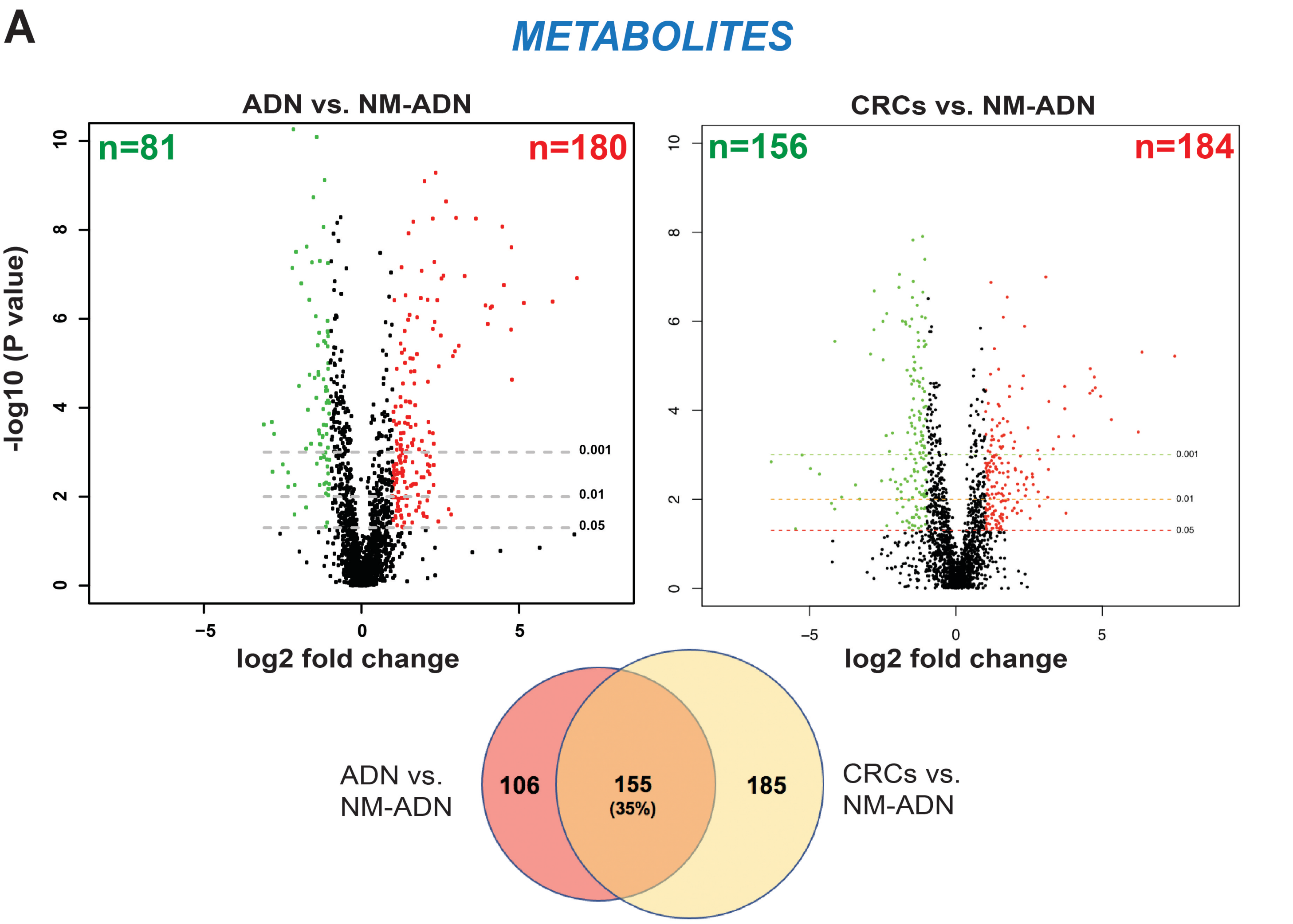


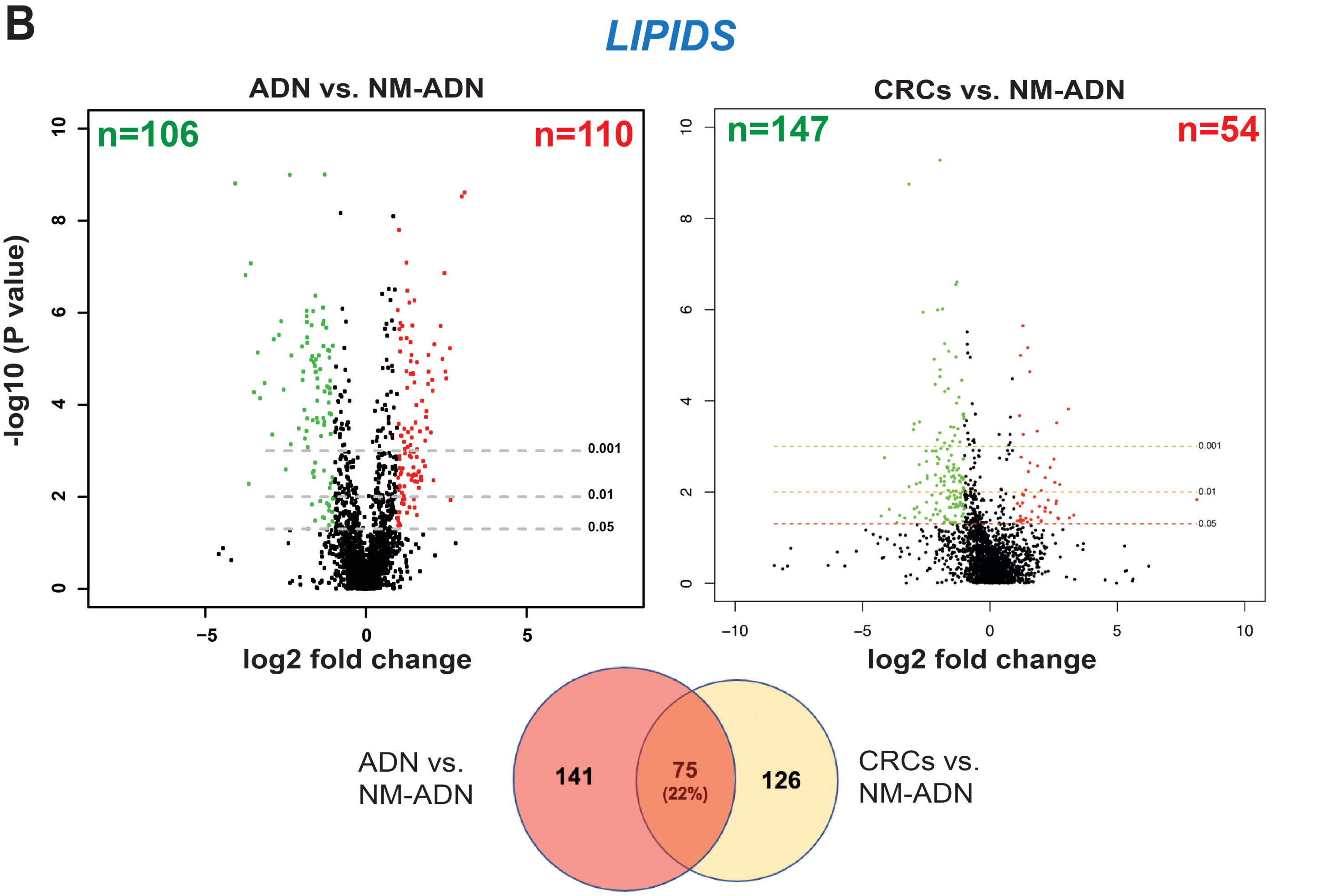
