## Supplementary Table 1 for "The metabolome and lipidome of colorectal adenomas and cancers"

### Supplementary Table 1. Internal standards mixture.

| Name | Label | Mass | Concentration* |
|---|---|---|---|
| **Metabolites:** | | | |
| Isovaleric Acid-1-13C, 99% 13C | 1 C13 | 103.03505 | 2 mM |
| SODIUM PYRUVATE-2-13C, 99% 13C | 1 C13 | 89.0194 | 2 mM |
| Malonic Acid-13C3, 99% 13C | All C13 | 107.021025 | 2 mM |
| Taurine-15N | 1 N15 | 126.011701 | 2 mM |
| D-GLUCOSE-1-13C, 99% 13C | 1 C13 | 181.066745 | 2 mM |
| D-Fructose-1-13C | 1 C13 | 181.066745 | 2 mM |
| AMP-13C10,15N5, 98% 13C, 96-98% 15N | 98% 13C, 96-98% 15N | 362.081813 | 2 mM |
| Dimethyl succinate-2,2,3,3-d4 | 4 D | 150.083018 | 2 mM |
| Sodium L-Lactate-1-13C solution, 99% 13C | 1 C13 | 91.03505 | 2 mM |
| D-Sorbitol-1-13C | 1 C13 | 183.082395 | 2 mM |
| UREA-15N2, 98% 15N | All N15 | 62.026433 | 2 mM |
| Algal Amino Acid Mixture-13C-15N: | 98% 13C, 98% 15N | | 2 mg/ml |
| **Lipids:** | | | |
| 1,2-diheptadecanoyl-*sn*-glycero-3-phospho-(1'-*rac*-glycerol) | | 772.523 | 1 mg/ml |
| 1,2-diheptadecanoyl-*sn*-glycero-3-phosphoethanolamine | | 719.547 | 1 mg/ml |
| 1-heptadecanoyl-2-hydroxy-*sn*-glycero-3-phosphocholine | | 509.348 | 1 mg/ml |
| 1,2-diheptadecanoyl-*sn*-glycero-3-phosphocholine | | 761.593 | 1 mg/ml |
| 1,2,3-triheptadecanoyl-glycerol | | 848.78 | 1 mg/ml |
| Margaric acid | | 270.2559 | 1 mg/ml |

* Concentration in the mixture added to the samples prior to LC_MS
