## Supplementary Table 2 for "The metabolome and lipidome of colorectal adenomas and cancers"

### Supplementary Table 2. The two LC-MS methods used in the study.

| Method Conditions | Metabolome | | | Lipidome | | |
|---|---|---|---|---|---|---|
| Column material | BEH Amide (Waters) | | | HSS T3 (Waters) | | |
| Particle Diameter | 1.8 um | | | 1.7 um | | |
| Column length | 15 cm | | | 5 cm | | |
| Column diameter | 200 um | | | 200 um | | |
| Injection solvent (1:5 diluted w/ analyte) | 90% acetonitrile, 9% methanol, 1% water, 50 mM ammonium acetate | | | Solvent A | | |
| Solvent A: | 0.5 mM ammonium acetate, pH 9 | | | 5 mM ammonium formate | | |
| Solvent B: | 95% acetonitrile, 0.5 mM ammonium acetate, pH 9 | | | 90% isopropanol, 10% acetonitrile, 5 mM ammonium formate | | |
| ESI polarity | negative | | | positive and negative | | |
| Gradient conditions: | Minutes | Composition B [%] | Flow rate [ul/min] | Minutes | Composition B [%] | Flow rate [ul/min] |
| | 0.00 | 90.0 | 6.0 | 0.00 | 0 | 4 |
| | 10.00 | 50.0 | 4.0 | 10.00 | 100 | 3 |
| | 12.00 | 50.0 | 4.0 | 15.00 | 100 | 3 |
| | 12.01 | 90.0 | 4.0 | 15.01 | 0 | 3 |
| | 14.00 | 90.0 | 4.0 | 20.00 | 0 | 3 |
