## Supplementary Table 3 for "The metabolome and lipidome of colorectal adenomas and cancers"

**Supplementary Table 3. Custom library of 190 core metabolites.** KEGG identifiers, molecular formulas, exact masses, partition coefficients, estimated limits of detection for the two applied LC methods are reported.

| Metabolite Information | | | | | HILIC (BEH Amide) | | Reversed Phase (HSS T3) | |
|---|---|---|---|---|---|---|---|---|
| **Name** | **KEGG ID** | **Formula** | **Mass** | **logP** | **LOD** | **Rf** | **LOD** | **Rf** |
| 1-Methylnicotinamide | C02918 | C7H9N2O | 137.0715 | -3.72 | | | 200 fmol | 1.2 |
| 1.3-Diaminopropane | C00986 | C3H10N2 | 74.0844 | -1 | | | 20 pmol | 3.4 |
| 2-Aminoethylphosphonate | C03557 | C2H8NO3P | 125.0242 | -1.55 | 20 pmol | 4.7 | 20 pmol | 1.1 |
| 2-Deoxy-D-ribose 1-phosphate | C00672 | C5H11O7P | 214.0242 | -2.14 | 2 pmol | 3.9 | | |
| 2-Oxobutanoate | C00109 | C4H6O3 | 102.0317 | 0.07 | 20 pmol | 1.1 | 2 pmol | 5.1 |
| 2-Oxoglutarate | C00026 | C5H6O5 | 146.0215 | -0.6 | 20 fmol | 2.9 | | |
| 2-Phenylacetamide | C02505 | C8H9NO | 135.0684 | 0.64 | 2 pmol | 1.1 | 20 fmol | 2.6 |
| 2-Phospho-D-glycerate | C00631 | C3H7O7P | 185.9929 | -2.24 | 20 pmol | 4.5 | 20 pmol | 1.0 |
| 2.3-Bisphospho-D-glycerate | C01159 | C3H8O10P2 | 265.9593 | -1.55 | | | 2 pmol | 0.9 |
| 3-Aminoisobutyric acid | C05145 | C4H9NO2 | 103.0633 | -2.97 | 2 pmol | 3.0 | 2 pmol | 1.1 |
| 3-Aminopropiononitrile | C05670 | C3H6N2 | 70.0531 | -0.9 | 20 pmol | 1.1 | 20 pmol | 1.0 |
| 3-Hydroxy-L-kynurenine | C03227 | C10H12N2O4 | 224.0797 | -2.09 | 2 pmol | 2.5 | 20 pmol | 1.1 |
| 3-Hydroxybutanoate | C01089 | C4H8O3 | 104.0473 | -0.5 | 2 pmol | 2.0 | 0.2 fmol | 5.1 |
| 3-Methoxy-4-hydroxymandelate | C05584 | C9H10O5 | 198.0528 | 0 | 200 fmol | 1.0 | | |
| 3-Methoxytyramine | C05587 | C9H13NO2 | 167.0946 | -0.04 | 20 pmol | 1.2 | 2 pmol | 2.1 |
| 3-Ureidopropionate | C02642 | C4H8N2O3 | 132.0535 | -0.98 | 2 pmol | 2.5 | | |
| 3.4-Dihydroxy-L-phenylalanine | C00355 | C9H11NO4 | 197.0688 | -2.32 | 20 pmol | 2.7 | 2 pmol | 1.2 |
| 3.4-Dihydroxymandelate | C05580 | C8H8O5 | 184.0372 | 0.72 | 20 pmol | 1.7 | 2 pmol | 0.8 |
| 4-Aminobutanoate | C00334 | C4H9NO2 | 103.0633 | -2.99 | 2 pmol | 3.5 | 2 pmol | 1.0 |
| 4-Pyridoxate | C00847 | C8H9NO4 | 183.0532 | -0.08 | 2 fmol | 1.0 | 2 fmol | 2.2 |
| 5-Amino-4-imidazolecarboxyamide | C04051 | C4H6N4O | 126.0542 | -1.16 | 2 pmol | 1.0 | 20 pmol | 5.6 |
| 5-Diphosphomevalonate | C01143 | C6H14O10P2 | 308.0062 | -1.17 | 2 pmol | 4.7 | 20 pmol | 0.9 |
| 5-Formyltetrahydrofolate | C03479 | C20H23N7O7 | 473.1659 | -0.46 | 0.2 fmol | 3.5 | 0.2 fmol | 2.3 |
| 5-Hydroxyindoleacetate | C05635 | C10H9NO3 | 191.0582 | 1.28 | | | | |
| 5-Methoxyindoleacetate | C05660 | C11H11NO3 | 205.0739 | 1.76 | | | | |
| 5-Methyltetrahydrofolate | C00440 | C20H25N7O6 | 459.1866 | -1.26 | 200 fmol | 3.9 | 20 fmol | 2.3 |
| 5-Phospho-alpha-D-ribose 1-diphos[illegible] | C00119 | C5H13O14P3 | 389.9518 | -6 | 20 pmol | 3.7 | 20 pmol | 0.9 |
| 5-Phosphomevalonate | C01107 | C6H13O7P | 228.0399 | -2.18 | 2 pmol | 4.0 | 20 pmol | 1.1 |
| 6-Deoxy-L-galactose | C01019 | C6H12O5 | 164.0685 | -2.39 | 20 pmol | 1.8 | | |
| Acetoacetate | C00164 | C4H6O3 | 102.0317 | -0.47 | 20 pmol | 1.8 | 20 pmol | 1.1 |
| Acetyl phosphate | C00227 | C2H5O5P | 139.9875 | -0.92 | 2 pmol | 3.5 | | |
| Acetyl-CoA | C00024 | C23H38N7O17P3S | 809.1258 | -0.58 | 2 fmol | 3.8 | 200 fmol | 2.3 |
| Adenine | C00147 | C5H5N5 | 135.0545 | -0.38 | 20 pmol | 4.5 | 2 pmol | 1.4 |
| Adenosine | C00212 | C10H13N5O4 | 267.0968 | -1.21 | 200 fmol | 1.1 | 20 fmol | 2.3 |
| Adenosine 3.5-bisphosphate | C00054 | C10H15N5O10P2 | 427.0294 | -1.63 | 0.2 fmol | 4.5 | 2 pmol | 1.3 |
| Adenylyl sulfate | C00224 | C10H14N5O10PS | 427.0199 | -1.64 | 20 fmol | 2.8 | | |
| ADP | C00008 | C10H15N5O10P2 | 427.0294 | -1.65 | 20 pmol | 4.9 | 2 pmol | 1.0 |
| ADP-ribose | C00301 | C15H23N5O14P2 | 559.0717 | -1.8 | 20 fmol | 3.7 | 200 fmol | 1.0 |
| Adrenaline | C00788 | C9H13NO3 | 183.0895 | -0.82 | 20 pmol | 3.8 | | |
| Agmatine | C00179 | C5H14N4 | 130.1218 | -1.01 | | | | |
| β-Alanine | C00099 | C3H7NO2 | 89.0477 | -3.26 | 2 pmol | 3.3 | 20 pmol | 1.0 |
| Alanine | C00041 | C3H7NO2 | 89.0477 | -3.05 | 2 pmol | 2.9 | 20 pmol | 1.0 |
| Allantoate | C00499 | C4H8N4O4 | 176.0546 | -2.12 | 200 fmol | 2.8 | 0.2 fmol | 4.7 |
| Allantoin | C02350 | C4H6N4O3 | 158.044 | -1.95 | 200 fmol | 1.3 | 20 pmol | 2.9 |
| Arabinose | C00259 | C5H10O5 | 150.0528 | -1.9 | 2 pmol | 1.9 | | |
| Arabitol | C00532 | C5H12O5 | 152.0685 | -3.2 | 2 pmol | 2.2 | 2 pmol | 5.1 |
| Ascorbate | C00072 | C6H8O6 | 176.0321 | -0.5 | 20 pmol | 2.6 | 0.2 fmol | 4.7 |
| Asparagine | C00152 | C4H8N2O3 | 132.0535 | -3.36 | 200 fmol | 3.3 | 2 pmol | 1.1 |
| Aspartate | C00402 | C4H7NO4 | 133.0375 | -3.52 | 2 fmol | 3.5 | 200 fmol | 0.9 |
| ATP | C00002 | C10H16N5O13P3 | 506.9957 | -0.84 | 200 fmol | 4.6 | 20 pmol | 12.4 |
| Biocytin | C05552 | C16H28N4O4S | 372.1831 | -0.69 | 20 fmol | 2.9 | 20 fmol | 2.3 |
| Biotin | C00120 | C10H16N2O3S | 244.0882 | 0.39 | | | 20 pmol | 1.0 |
| Butanoyl-CoA | C00136 | C25H42N7O17P3S | 837.1571 | -0.22 | 200 fmol | 3.4 | 2 pmol | 2.5 |
| Cadaverine | C01672 | C5H14N2 | 102.1157 | -0.27 | | | 20 pmol | 1.7 |
| cis-Aconitate | C00417 | C6H6O6 | 174.0164 | -0.41 | 20 fmol | 3.9 | 0.2 fmol | 1.4 |
| Citrate | C00158 | C6H8O7 | 192.027 | -1.33 | 2 pmol | 4.8 | 200 fmol | 1.0 |
| CoA | C00010 | C21H36N7O16P3S | 767.1152 | -0.61 | 20 pmol | 3.9 | 200 fmol | 2.2 |
| Coniferyl alcohol | C00590 | C10H12O3 | 180.0786 | 1.51 | 2 pmol | 1.1 | 20 pmol | 2.9 |
| Cysteine | C00097 | C3H7NO2S | 121.0197 | -2.57 | 2 pmol | 3.9 | 20 pmol | 1.0 |
| dADP | C00206 | C10H15N5O9P2 | 411.0345 | -1.57 | 200 fmol | 4.2 | 2 pmol | 1.4 |
| dAMP | C00360 | C10H14N5O6P | 331.0682 | -2.44 | 2 fmol | 3.7 | 2 pmol | 1.7 |
| Decanoyl-CoA | C05274 | C31H54N7O17P3S | 921.251 | 0.87 | 20 fmol | 2.9 | 2 pmol | 4.8 |
| Deoxyadenosine | C00559 | C10H13N5O3 | 251.1018 | -0.95 | 200 fmol | 1.1 | 20 fmol | 2.3 |
| Deoxyguanosine | C00330 | C10H13N5O4 | 267.0968 | -1.75 | 2 fmol | 1.2 | 200 fmol | 2.2 |
| dGDP | C00361 | C10H15N5O10P2 | 427.0294 | -1.45 | 20 pmol | 4.9 | 2 pmol | 1.0 |
| dGMP | C00362 | C10H14N5O7P | 347.0631 | -1.87 | 0.2 fmol | 4.1 | 2 pmol | 1.2 |
| dGTP | C00286 | C10H16N5O13P3 | 506.9957 | -0.61 | 0.2 fmol | 5.1 | 2 pmol | 1.0 |
| Dihydrofolate | C00415 | C19H21N7O6 | 443.1553 | -3.2 | | | | |
| Dihydroorotate | C00337 | C5H6N2O4 | 158.0328 | -1.7 | | | 20 pmol | 4.4 |
| Dimethylallyl diphosphate | C00235 | C5H12O7P2 | 246.0058 | 0.3 | 0.2 fmol | 3.9 | | |

| | | | | | | | | |
|---|---|---|---|---|---|---|---|---|
| Dopamine | C03758 | C8H11NO2 | 153.079 | -0.4 | 20 pmol | 1.3 | 20 pmol | 2.2 |
| Ethanolamine phosphate | C00346 | C2H8NO4P | 141.0191 | -2.5 | 2 pmol | 4.5 | 200 fmol | 1.0 |
| FMN | C00061 | C17H21N4O9P | 456.1046 | -0.78 | 200 fmol | 3.5 | 200 fmol | 2.5 |
| Fructose | C00095 | C6H12O6 | 180.0634 | -2.4 | 20 pmol | 2.9 | 20 pmol | 1.4 |
| Fructose 1.6-bisphosphate | C05378 | C6H14O12P2 | 339.996 | -1.53 | 2 pmol | 4.8 | 20 pmol | 0.9 |
| Fructose 6-phosphate | C05345 | C6H13O9P | 260.0297 | -2.11 | 0.2 fmol | 4.2 | 20 pmol | 0.9 |
| Fucose 1-phosphate | C02985 | C6H13O8P | 244.0348 | -1.53 | 20 fmol | 4.1 | | |
| Fumarate | C00122 | C4H4O4 | 116.011 | 0.21 | 2 pmol | 2.9 | 20 pmol | 3.4 |
| Galactose | C00124 | C6H12O6 | 180.0634 | -2.57 | 2 pmol | 2.6 | 20 pmol | 2.2 |
| Galactose 1-phosphate | C00446 | C6H13O9P | 260.0297 | -2 | 0.2 fmol | 4.3 | | |
| GDP | C00035 | C10H15N5O11P2 | 443.0243 | -1.51 | 2 pmol | 5.3 | 200 fmol | 0.9 |
| GDP-L-fucose | C00325 | C16H25N5O15P2 | 589.0822 | -1.69 | 2 fmol | 4.1 | 200 fmol | 1.0 |
| GDP-mannose | C00096 | C16H25N5O16P2 | 605.0772 | -1.76 | 2 fmol | 4.4 | 2 pmol | 1.0 |
| Gentisate aldehyde | C05585 | C7H6O3 | 138.0317 | 1.02 | 20 fmol | 1.0 | 20 pmol | 2.8 |
| Glucarate | C00818 | C6H10O8 | 210.0376 | -3.6 | 200 fmol | 3.8 | 2 pmol | 0.9 |
| Gluconic acid | C00257 | C6H12O7 | 196.0583 | -1.87 | 200 fmol | 3.1 | 20 pmol | 5.1 |
| Glucosamine 6-phosphate | C00352 | C6H14NO8P | 259.0457 | -2.6 | 20 pmol | 4.6 | 2 pmol | 1.0 |
| Glucose | C00031 | C6H12O6 | 180.0634 | -2.57 | 2 pmol | 2.6 | 20 pmol | 1.0 |
| Glucose 6-phosphate | C00668 | C6H13O9P | 260.0297 | -2.06 | 0.2 fmol | 4.4 | 2 pmol | 2.8 |
| Glucuronate | C00191 | C6H10O7 | 194.0427 | -2 | 20 pmol | 3.5 | | |
| Glutamate | C00025 | C5H9NO4 | 147.0532 | -3.54 | 20 fmol | 3.3 | 20 pmol | 1.1 |
| Glutamine | C00064 | C5H10N2O3 | 146.0691 | -3.32 | 200 fmol | 3.3 | 0.2 fmol | 5.1 |
| Glutaryl-CoA | C00527 | C26H42N7O19P3S | 881.1469 | -0.48 | 20 fmol | 4.3 | 20 fmol | 2.3 |
| Glyceraldehyde 3-phosphate | C00118 | C3H7O6P | 169.998 | -1.69 | 2 pmol | 4.2 | 20 pmol | 1.0 |
| Glycerate | C00258 | C3H6O4 | 106.0266 | -1.72 | 20 fmol | 2.4 | 20 pmol | 1.3 |
| Glycerol 3-phosphate | C00093 | C3H9O6P | 172.0137 | -1.84 | 200 fmol | 3.8 | 2 pmol | 0.9 |
| Glycine | C00037 | C2H5NO2 | 75.032 | -3.34 | 20 pmol | 3.3 | 20 pmol | 1.0 |
| Glyoxylate | C00048 | C2H2O3 | 74.0004 | -0.59 | 2 pmol | 2.4 | | |
| GTP | C00044 | C10H16N5O14P3 | 522.9907 | -0.63 | 200 fmol | 5.1 | 2 pmol | 0.9 |
| Guanine | C00242 | C5H5N5O | 151.0494 | -0.9 | 20 pmol | 3.2 | 200 fmol | 1.2 |
| Guanosine | C00387 | C10H13N5O5 | 283.0917 | -2.06 | 200 fmol | 1.9 | 20 pmol | 1.8 |
| Hexanoyl-CoA | C05270 | C27H46N7O17P3S | 865.1884 | 0.07 | 2 fmol | 3.2 | 200 fmol | 3.1 |
| Homogentisate | C00544 | C8H8O4 | 168.0423 | 0.81 | 2 pmol | 1.0 | 0.2 fmol | 5.1 |
| Homovanillate | C05582 | C9H10O4 | 182.0579 | 1.02 | 2 pmol | 1.0 | 2 pmol | 13.2 |
| Hypoxanthine | C00262 | C5H4N4O | 136.0385 | -0.55 | 0.2 fmol | 1.1 | 200 fmol | 1.4 |
| IDP | C00104 | C10H14N4O11P2 | 428.0134 | -2.47 | 20 pmol | 4.7 | 2 pmol | 1.0 |
| IMP | C00130 | C10H16N5O12P3 | 491.0008 | -0.66 | 20 pmol | 4.7 | | |
| Indole-3-acetate | C00954 | C10H9NO2 | 175.0633 | 1.87 | 200 fmol | 1.1 | 200 fmol | 2.8 |
| Inosine | C00294 | C10H12N4O5 | 268.0808 | -1.87 | 20 fmol | 1.2 | 20 pmol | 1.8 |
| Inositol | C00137 | C6H12O6 | 180.0634 | -2.08 | 200 fmol | 4.0 | | |
| Isocitrate | C00311 | C6H8O7 | 192.027 | -0.35 | 2 pmol | 4.4 | 2 pmol | 0.9 |
| Isopentenyl diphosphate | C00129 | C5H12O7P2 | 246.0058 | 0.04 | 200 fmol | 3.9 | 2 pmol | 2.9 |
| ITP | C00081 | C10H15N4O14P3 | 507.9798 | -0.67 | 20 pmol | 5.2 | 200 fmol | 0.9 |
| Lactose | C00243 | C12H22O11 | 342.1162 | -5.03 | 20 fmol | 4.0 | 0.2 fmol | 5.8 |
| Lauroyl-CoA | C01832 | C33H58N7O17P3S | 949.2823 | 1.35 | 20 fmol | 2.8 | 2 pmol | 5.2 |
| Malate | C00149 | C4H6O5 | 134.0215 | -0.87 | 0.2 fmol | 3.5 | | |
| Malonate | C00383 | C3H4O4 | 104.011 | -0.6 | 20 fmol | 3.4 | 200 fmol | 13.2 |
| Malonyl-CoA | C00083 | C24H38N7O19P3S | 853.1156 | -0.62 | 200 fmol | 4.3 | 2 pmol | 2.2 |
| Maltose | C00208 | C12H22O11 | 342.1162 | -5.03 | 0.2 fmol | 3.8 | 2 fmol | 5.8 |
| Mannitol | C00392 | C6H14O6 | 182.079 | -2.68 | 20 fmol | 2.6 | 2 pmol | 2.6 |
| Mannose | C00159 | C6H12O6 | 180.0634 | -2.57 | 200 fmol | 4.0 | 20 pmol | 13.2 |
| Mannose 1-phosphate | C00636 | C6H13O9P | 260.0297 | -2 | 0.2 fmol | 4.3 | 2 pmol | 1.0 |
| Mannose 6-phosphate | C00275 | C6H13O9P | 260.0297 | -2.06 | 200 fmol | 4.3 | 20 pmol | 1.1 |
| Melatonin | C01598 | C13H16N2O2 | 232.1212 | 1.42 | 200 fmol | 1.0 | 20 fmol | 3.2 |
| Methylglyoxal | C00546 | C3H4O2 | 72.0211 | -0.38 | 200 fmol | 1.8 | | |
| Methylmalonate | C02170 | C4H6O4 | 118.0266 | 0.17 | 20 pmol | 3.0 | 0.2 fmol | 5.1 |
| N-(L-Arginino)succinate | C03406 | C10H18N4O6 | 290.1226 | -3.25 | 20 fmol | 4.5 | 200 fmol | 1.0 |
| N-Acetyl-D-glucosamine | C00140 | C8H15NO6 | 221.0899 | -2.6 | 200 fmol | 2.0 | 2 pmol | 1.0 |
| N-Acetyl-L-aspartate | C01042 | C6H9NO5 | 175.0481 | -0.79 | 2 fmol | 3.3 | 2 pmol | 0.9 |
| N-Acetylneuraminate | C00270 | C11H19NO9 | 309.106 | -2.78 | 2 fmol | 3.1 | 2 pmol | 1.0 |
| N-Acetylputrescine | C02714 | C6H14N2O | 130.1106 | -0.84 | | | 2 pmol | 1.2 |
| N-Acetylserotonin | C00978 | C12H14N2O2 | 218.1055 | 0.98 | 200 fmol | 1.0 | 200 fmol | 2.5 |
| N-Methylhistamine | C05127 | C6H11N3 | 125.0953 | -0.57 | 20 pmol | 3.6 | 20 pmol | 1.4 |
| N-Methylserotonin | C06212 | C11H14N2O | 190.1106 | 1.55 | 20 pmol | 1.3 | 200 fmol | 2.2 |
| N-Methyltryptamine | C06213 | C11H14N2 | 174.1157 | 2.02 | 20 pmol | 1.2 | 200 fmol | 2.5 |
| NAD+ | C00003 | C21H28N7O14P2 | 663.1091 | -1.18 | 2 pmol | 4.0 | | |
| NADH | C00004 | C21H29N7O14P2 | 665.1248 | -1.45 | 20 fmol | 3.4 | 2 pmol | 1.4 |
| Noradrenaline | C00547 | C8H11NO3 | 169.0739 | -1.4 | 20 pmol | 2.9 | 20 pmol | 5.6 |
| Normetanephrine | C05589 | C9H13NO3 | 183.0895 | -0.71 | 2 pmol | 1.2 | 2 pmol | 1.1 |
| Octanoyl-CoA | C01944 | C29H50N7O17P3S | 893.2197 | -3 | 2 pmol | 3.1 | 2 pmol | 4.1 |
| Orotate | C00295 | C5H4N2O4 | 156.0171 | -0.89 | 20 fmol | 1.6 | 20 pmol | 4.5 |
| Palmitoyl-CoA | C00154 | C37H66N7O17P3S | 1005.3449 | 2.35 | 200 fmol | 2.8 | 20 pmol | 6.0 |
| Phenethylamine | C05332 | C8H11N | 121.0891 | 1.41 | | | 200 fmol | 2.3 |
| Phenylacetaldehyde | C00601 | C8H8O | 120.0575 | 1.75 | 2 pmol | 1.0 | 0.2 fmol | 5.0 |
| Putrescine | C00134 | C4H12N2 | 88.1 | -0.98 | | | 20 pmol | 2.8 |
| Pyridoxal | C00250 | C8H9NO3 | 167.0582 | 0.02 | 20 fmol | 1.1 | 200 fmol | 1.4 |
| Pyridoxal phosphate | C00018 | C8H10NO6P | 247.0246 | -0.55 | 200 fmol | 3.9 | 200 fmol | 1.5 |

| | | | | | | | | |
|---|---|---|---|---|---|---|---|---|
| Pyridoxamine | C00534 | C8H12N2O2 | 168.0899 | -1.23 | 200 fmol | 2.4 | 2 pmol | 1.2 |
| Pyruvate | C00022 | C3H4O3 | 88.016 | -0.38 | 20 pmol | 1.5 | 20 pmol | 1.0 |
| quinone | C00472 | C6H4O2 | 108.0211 | 0.21 | | | 20 pmol | 5.1 |
| Raffinose | C00492 | C18H32O16 | 504.169 | -3.36 | 2 fmol | 4.8 | 2 pmol | 1.0 |
| Ribitol | C00474 | C5H12O5 | 152.0685 | -2.53 | 2 pmol | 1.8 | | |
| Riboflavin | C00255 | C17H20N4O6 | 376.1383 | -1.05 | 200 fmol | 0.8 | 200 fmol | 0.9 |
| Ribose | C00121 | C5H10O5 | 150.0528 | -2.65 | 20 pmol | 1.3 | 20 pmol | 2.9 |
| Ribose 1-phosphate | C00620 | C5H11O8P | 230.0192 | -2.04 | 2 pmol | 4.0 | | |
| Ribose 5-phosphate | C00117 | C5H11O8P | 230.0192 | -2.07 | 2 pmol | 4.1 | | |
| Ribulose 5-phosphate | C00199 | C5H11O8P | 230.0192 | -2.07 | 200 fmol | 3.8 | 20 pmol | 1.2 |
| Selenate | C05697 | H2SeO4 | 145.9118 | | | | | |
| Serine | C00740 | C3H7NO3 | 105.0426 | -3.42 | 200 fmol | 3.2 | 20 pmol | 1.2 |
| Serine | C00065 | C3H7NO3 | 105.0426 | -3.42 | 200 fmol | 3.3 | 20 pmol | 1.0 |
| Sinapyl alcohol | C02325 | C11H14O4 | 210.0892 | 1.36 | 20 pmol | 3.0 | | |
| Sorbitol | C00794 | C6H14O6 | 182.079 | -2.68 | 20 fmol | 2.5 | 20 pmol | 1.0 |
| Sorbitol 6-phosphate | C01096 | C6H15O9P | 262.0454 | -2.32 | 200 fmol | 4.3 | 2 pmol | 0.9 |
| Spermidine | C00315 | C7H19N3 | 145.1579 | -0.62 | | | | |
| Spermine | C00750 | C10H26N4 | 202.2157 | -0.66 | | | | |
| Succinate | C00042 | C4H6O4 | 118.0266 | -0.53 | 2 pmol | 3.2 | 20 pmol | 2.8 |
| Succinate semialdehyde | C00232 | C4H6O3 | 102.0317 | -0.47 | 200 fmol | 2.8 | | |
| Succinyl-CoA | C00091 | C25H40N7O19P3S | 867.1313 | -6.7 | 200 fmol | 4.3 | 200 fmol | 2.3 |
| Sucrose | C00089 | C12H22O11 | 342.1162 | -2.63 | 0.2 fmol | 3.4 | 20 pmol | 1.0 |
| Sulfite | C00094 | H2SO3 | 81.9725 | -2.7 | 20 pmol | 1.1 | | |
| Taurine | C00245 | C2H7NO3S | 125.0147 | -2.19 | 200 fmol | 2.5 | 2 pmol | 1.1 |
| Taurocholate | C05122 | C26H45NO7S | 515.2917 | 0.79 | 20 fmol | 1.0 | 200 fmol | 4.6 |
| Tetradecanoyl-CoA | C02593 | C35H62N7O17P3S | 977.3136 | 1.84 | 20 fmol | 2.9 | 2 pmol | 5.6 |
| Tetrahydrofolate | C00101 | C19H23N7O6 | 445.171 | -1.45 | | | | |
| Thiamin diphosphate | C00068 | C12H19N4O7P2S | 425.045 | -0.1 | 20 pmol | 5.7 | 20 pmol | 1.4 |
| Thiamin monophosphate | C01081 | C12H18N4O4PS | 345.0786 | -1.68 | 20 pmol | 4.9 | 20 pmol | 1.1 |
| Thiamine | C00378 | C12H17N4OS | 265.1123 | -2.11 | 2 pmol | 2.2 | 200 fmol | 2.2 |
| Trehalose | C01083 | C12H22O11 | 342.1162 | -2.98 | 0.2 fmol | 3.9 | 0.2 fmol | 5.8 |
| Trypanothione disulfide | C03170 | C27H47N9O10S2 | 721.2887 | | 2 pmol | 6.5 | 20 pmol | 1.4 |
| Tryptamine | C00398 | C10H12N2 | 160.1 | 1.21 | | | | |
| Tryptophan | C00078 | C11H12N2O2 | 204.0899 | -1.1 | 200 fmol | 1.3 | 2 pmol | 2.3 |
| Tyramine | C00483 | C8H11NO | 137.0841 | -0.14 | 2 pmol | 1.2 | 20 pmol | 1.5 |
| Tyrosine | C00082 | C9H11NO3 | 181.0739 | -2.39 | 200 fmol | 2.5 | 2 pmol | 1.1 |
| UDP-D-galactose | C00052 | C15H24N2O17P2 | 566.055 | -6.5 | 2 fmol | 4.0 | 20 pmol | 0.9 |
| UDP-glucuronate | C00167 | C15H22N2O18P2 | 580.0343 | -1.21 | 2 fmol | 4.4 | 2 pmol | 0.9 |
| Urate | C00366 | C5H4N4O3 | 168.0283 | -1.12 | | | 0.2 fmol | 5.0 |
| Urea | C00086 | CH4N2O | 60.0324 | -1.78 | 0.2 fmol | 2.6 | | |
| Xanthine | C00385 | C5H4N4O2 | 152.0334 | -0.65 | 20 pmol | 1.1 | 20 pmol | 1.4 |
| Xylitol | C00379 | C5H12O5 | 152.0685 | -3.2 | 2 pmol | 2.3 | | |
| Xylose | C00181 | C5H10O5 | 150.0528 | -2.57 | 20 pmol | 3.9 | | |
